# Transcription factor 19 is an androgen responsive gene that modulates vessel homeostasis and sustains metastatic prostate cancer

**DOI:** 10.1101/2025.02.04.636232

**Authors:** Amaia Ercilla, Jana R. Crespo, Saioa Garcia-Longarte, Marta Fidalgo, Natalia Martin-Martin, Onintza Carlevaris, Ianire Astobiza, Sonia Fernández-Ruiz, Marc Guiu, Laura Bárcena, Isabel Mendizabal, Ana M. Aransay, Mariona Graupera, Roger R. Gomis, Arkaitz Carracedo

## Abstract

Prostate cancer is a prevalent tumor type that, despite being highly curable, progresses to metastatic disease in a fraction of patients, thus accounting for more than 350.000 annual deaths worldwide. In turn, uncovering the molecular insights of metastatic disease is instrumental to improve the survival rate of prostate cancer patients. By means of gene expression metanalysis in multiple prostate cancer patient cohorts, we identified a set of genes that are differentially expressed in aggressive prostate cancer. *Transcription factor 19 (TCF19)* stood out as an unprecedented epithelial gene upregulated in metastatic disease, with prognostic potential and associated with the activity of androgen receptor. By combining computational and empiric approaches, our data revealed that TCF19 is required for full metastatic capacity and its depletion influences core cancer-related processes, such as vascular permeability, supporting the role of this gene in the dissemination of prostate tumor cells.

## Introduction

Prostate cancer (PCa) is the second most common cancer type in male worldwide affecting millions of men every year. Despite being highly curative, due to its high incidence, PCa accounts for more than 350,000 annual deaths, making it one of the leading causes of cancer-associated deaths in men^1^. The incorporation of computational biology approaches over the last years has provided a more multidisciplinary view to PCa research, increasing our understanding of the molecular basis of this disease. For instance, pan-cancer whole genome sequencing studies have shown that, in contrast to other tumor types, the genomic landscape of advanced disease differs profoundly from the indolent tumors^2^. Despite these advances, the 5-year survival rate of patients that suffer the most aggressive form of metastatic PCa remains below 30-40%, and the key determinants of its aggressiveness still need to be elucidated^1^.

Androgen receptor (AR) plays a central role in the differentiation and homeostasis of the prostate tissue. Although it is commonly considered a transcription activator, increasing evidence supports an additional role of AR as a transcriptional repressor^3^. AR activity is commonly hijacked by tumor cells for continuous growing, underscoring the relevance of targeting this receptor in prostate cancer. However, although androgen deprivation therapy initially shows a favorable response, resistance eventually arises, leading to castration-resistant prostate cancer (CRPC)^1^.

Transcription factor 19 (TCF19) was originally identified as a serum-regulated factor expressed at the G1/S boundary and S phase^4^. It harbors a forkhead-associated (FHA) domain and a plant homeodomain (PHD) finger through which binds to histone 3 lysine 4 trimethylation^5^. Genome-wide association studies postulated a role of TCF19 in type I and II diabetes^6–8^, and more recent studies have reported a role of TCF19 in glucose metabolism^9–11^, including the recruitment of nucleosome-remodeling-deacetylase (NuRD) complex to the promoters of genes involved in *de novo* glucose production^5^. Besides controlling the recruitment of NuRD, TCF19 can form different transcription activation/repression complexes with p53 to regulate glycolysis and OXPHOS pathways^12^.

During the past years, increasing evidence support the contribution of TCF19 to tumor progression by controlling cellular proliferation in several cancer types^13–20^through a mechanism that is mediated by targeting FOXO1 pathway^14,16^. Very recently, roles of TCF19 that go beyond promoting cellular proliferation such as inducing epithelial to mesenchymal (EMT) transition^21^ or facilitating CD8+ T cell exhaustion^22^ have also been reported. Despite these recent advances in the role of TCF19, its contribution to prostate cancer progression and in particular to metastatic disease have never been addressed before.

In this work, taking advantage of PCa transcriptomics data placed in public repositories, we designed an integrative multi-dataset metanalysis to uncover novel drivers of aggressive PCa. Using a stringent criterion, we identified 9 genes whose expression 1) was consistently deregulated in metastatic specimens and 2) exhibited prognostic potential. *TCF19* stood out among the most robust epithelial gene fulfilling these criteria, and we proceeded to study its regulation and the consequences of its perturbation in prostate tumor cells.

## Materials and methods

### Gene expression Data Metanalysis

The metanalysis was performed with available information downloaded from Gene Expression Omnibus (GEO) with the exception of TCGA dataset, which was obtained from (https://gdac.broadinstitute.org/). The pre-processed dataset normalization was reviewed and corrected when required. A background correction, Log_2_ normalization and quartile normalization was applied when needed. In the Cox plots, the optimal degree of freedom was determined based on the Akaike Information Criterion (AIC) to fit smooth curves to the data for each gene and dataset. A non-parametric Spearman correlation test was applied to analyze the association between two genes or signatures. Patient data-containing public datasets were stratified based on the mean *TCF19* levels to perform Gene Set Enrichment analysis (GSEA). The data from our RNA-seq analysis was stratified based on shScramble or shTCF19 expression for GSEA^23^ and xCell^24^ analysis.

### Cell culture and treatments

Human prostate cancer PC3, DU145 and LNCaP cells were purchased at Leibniz Institut DSMZ (Deutsche Sammlung con Mikroorganismen und Zelkulturen GmbH), with the corresponding certificate of authenticity. Human prostate cancer cell lines 22RV1 and VCaP were purchased from American Type Culture Collection, who provided authentication certificate. Human prostate cancer cell C4-2 cells were a kind gift provided by the laboratory of Dr. Pier Paolo Pandolfi. Cell lines were periodically subjected to microsatellite-based identity validation. Cell lines were tested for mycoplasma contamination routinely using MycoAlert detection Kit (Lonza; LT07-318). PC3, DU145 and VCaP cells were cultured in Dulbecco’s modified eagle medium (DMEM) (Gibco™; 41966-029). LNCaP, 22RV1 and VCaP cells were maintained in Roswell park memorial institute (RPMI 1640; 11875-093) medium. Both culture mediums were supplemented with 10% of inactivated fetal bovine serum (FBS) (Gibco™; 10270-106) and 1% of penicillin/ streptomycin (Gibco™; 15140-122).

### Lentiviral production and cell line generation

HEK293FT cells were used for lentiviral production. Lentiviral vectors expressing short hairpins (shRNAs) against human Scramble and TCF19 were purchased from Sigma-Aldrich. Cells were transfected with lentiviral vectors following standard procedures. Puromycin (2 μg/ml; Sigma-Aldrich; P8833) was used for selection. The shRNA sequences are detailed in Table S1.

### Cellular assays

#### Proliferation assays

5000 cells/well were seeded in 12-well plates on triplicate. Cells were fixed with 10% formalin (Avantor) after 0, 2 and 4 days and stained with crystal violet (0.1% in 20% methanol; Sigma-Aldrich) for 30 min. Plates were air dried at least overnight before dissolving the crystal violet in 10% acetic acid for reading the absorbance at 595.

#### Cell cycle analysis

For cell cycle analysis, cells were seeded in black-wall 96-well plates (BIO-Greiner) and labelled with 10 µM 5-etinil-2’-desoxiuridina (EdU; Invitrogen; A10044) for 30min before fixing them with 4% formaldehyde. For EdU detection, cells were permeabilized with 0.2% triton-containing PBS for 30min and click reaction was performed by incubating the cells for 1h at RT on click-IT buffer (100 mM Tris-HCl pH 8, 2 mM CuSO4 (Sigma-Aldrich), 1 ng Alexa Fluor 488 Azide (Life Technologies; A10266) and 100 mM ascorbic acid (Sigma-Aldrich; PHR1008-2G)). Mitotic cells were detected by immunostaining with Phospho-Ser/Thr-Pro MPM-2 (Sigma-Aldrich; 05-368) antibody. DNA was counterstained with Hoechst (Life technologies, H3570).

#### Foci and colony formation assays

The ability to grow individualized was measured by seeding 250 cells/well in a 6-well plate on triplicate. Cells were fixed with 10% formalin (Avantor) and stained with 0.1% crystal violet (Sigma-Aldrich) in 20% methanol after 7 (DU145) or 15 (PC3, 22RV1) days. The number of foci was counted using Fiji software.

Anchorage-independent growth was measured by seeding 2500 (PC3) or 5000 (DU145, 22RV1) cells on agar-coated 6-well dishes on triplicate as previously described^25^. Colonies were imaged using an Olympus IX-83 inverted microscope operated by CellSens software and the number of visible colonies was counted using Fiji software.

#### Wound Healing assays

Cell migration rate was measured as previously described^25^. Briefly, cells were seeded at high confluency on 6-well plates on triplicate, a wound was created with the help of a pipet tip and images were acquired at different timepoints to calculate the linear growth of wound closure. Fiji software was used to quantify the wounded area.

#### Invasive growth assays

Spheroid assays were performed by preparing 700 cell drops of 25 µL with 6% methylcellulose (Sigma-Aldrich; M0387) and incubating them at 37°C and 5% CO_2_ for 48 hours. Spheroids were collected and embedded in collagen I solution (Advanced BioMatrix PureCol; 5005). Pictures were taken at the indicated timepoints using an Olympus IX-83 inverted microscope operated by CellSens software. Invasive growth was calculated by the differential area between initial and final timepoint using FiJi software.

### In vivo assays

All mouse experiments were carried out following the ethical guidelines established by the Biosafety and Animal Welfare Committee at CIC bioGUNE (Spanish acronym for center for cooperative research in Biosciences). The procedures employed were carried out following the recommendations from the Association for Assessment and Accreditation of Laboratory Animal Care (AAALAC). Mice were maintained in a controlled environmental conditions, 361 with time-controlled lighting on standard 12:12 light:dark cycles, 30-50% of humidity and 362 controlled temperature at 22±2°C. Diet and water were provided *ad libitum*. Orthotopic xenotransplant assays were performed by injecting 7.5 x 10^5^ GFP-Luc expressing PC3 cells into the ventral prostate lobes of 10 Nu/Nu immunodeficient males. 1.5 x 10^5^ GFP-Luc expressing PC3 cells were injected in the left ventricle of 7 Nu/Nu immunodeficient males for intracardiac metastatic assays. The sh2 was used in both cases to generate TCF19-depleted cells. Tumor growth and dissemination were followed by measuring bioluminescence with IVIS technology (PerkinElmer) for the indicated days. Intra-orbital injections of 50 µL luciferase at 15 mg/mL were used during the follow-up.

### Real-time quantitative RT-PCR (qRT-PCR)

Maxwell RSC simplyRNA Cells Kit (Promega; AS1390) was used to isolate total RNA. Complementary DNA was produced using Maxima™ H Minus cDNA Synthesis Master Mix (Invitrogen; M1682) following manufacturers guidelines. qRT-PCR were run in a Viia7, QS5 or QS6 Real-Time PCR Systems (Applied Biosystems) using the Taqman probes from Life Technologies or the primer/probes from Universal Probe Library from Roche detailed in Table S2. The expression of individual genes was calculated and normalized to the indicated reference gene.

### Immunohistochemistry and immunofluorescence experiments

For the analysis of Ki67, P-pRb (S807/811) and Cleaved-Caspase 3 levels in the tumor sections of the orthotopic experiments, paraffin embedded tissues were deparaffinize, rehydrated, and antigen retrieval was performed using citrate pH=6 buffer following standard procedures. Tissue sections were permeabilized by incubation with 0.2% triton-containing PBS for 30min at RT and blocked with 2%FBS-1%BSA-containing PBS. For IHC, tissues were incubated with Ki67 (Abcam, ab16667) antibody overnight at 4°C and 1:100 dilution. Hematoxylin/eosin was applied as counterstaining. Secondary antibody and DAB staining were performed by routinely used procedures. For IF, primary antibodies against Phospho-Ser/Thr-Pro MPM-2 (Sigma-Aldrich; 05-368; 1:250), P-pRb S807/811-647 (Cell Signaling, #8974; 1:50) or Cleaved-Caspase 3 (Cell Signaling, #9661; 1:400) were incubated for 1h or 2h (in the case of Cleaved-Caspase 3), followed by a 1h incubation with Alexa Fluor (Thermo Fisher scientific; 1:500) secondary antibodies. DNA was counterstained with Hoechst 1:2000 (Life technologies, H3570).

For the analysis of CD31, Alpha Smooth Muscle Actin (αSMA), TER119, B-catenin and Desmin levels in the tumor sections of the orthotopic experiments, paraffin sections (5 µm) were incubated at 60°C for 1h, deparaffinized and rehydrated. Antigen retrieval was then performed with citrate buffer at 100°C for 10 minutes, followed by permeabilization (0,2% Tween-20 in PBS) at RT for 30 minutes. Sections were blocked for 30 minutes with 10% horse serum (HS), 2% BSA in PBS and subsequentially incubated with the primary antibodies (antibodies were diluted in 5% HS, 1%BSA, in PBS) overnight at 4°C. The following primary antibodies were used in each case: CD31 (Abcam, ab28364), αSMA (Sigma-Aldrich, C6198, Cy3 conjugated), TER119 (R&D systems, MAB1125), B-catenin (BD Transduction Laboratories, 610154) and Desmin (Abcam, ab195177, AF647 conjugated). Alexa-Fluor secondary antibodies from Invitrogen (diluted the same way as primary antibodies) were added for 2h at RT and slides were mounted using Immu-Mount (Epredia, 9990402). In between steps slides were washed 3 times with PBS for 5 minutes, at RT, in movement. DNA was counterstained using DAPI (Invitrogen, S33025). Images were acquired using a Leica Stellaris 8 confocal microscope. All tile scans were acquired at 10x magnification and images for quantification were acquired at 20x or 40x magnification. Analysis and quantification of the images was done with the Imaris software (version 10.1.1).

### RNA-Seq analysis

Total RNA was extracted using Maxwell RSC simplyRNA Cells Kit (Promega; AS1390) and Total RNA libraries were prepared at the Genomics platform of CIC bioGUNE using TruSeq Stranded Total RNA with Ribo-Zero Globin kit (Illumina Inc., Cat.# 20020612) and TruSeq RNA CD Index Plate (Illumina Inc., Cat.# 20019792) following “TruSeq Stranded Total RNA Sample Prep-guide (Part # 15031048 Rev. E)”. Libraries were sequenced on an Illumina NovaSeq 6000 instrument to generate at least 40 million paired-end 100 bp reads.

Reads were aligned to the human reference genome (hg38) using STAR (version_2.7.5c) in two-pass mode following STAR best practices and recommendations^26^. The quality of the data was evaluated using STAR (version 2.7.5c)^26^ and samtools (version 1.15)^27^. PCR duplicates were removed from aligned bam files using samtools (version 1.15)^27^. Read counts were extracted from the aligned bam files using subread’s FeatureCounts (version 2.0.3)^28^. Normalization of read counts for analysis was done using the Ratio of the Variance method which accounts for inter-sample variance and the differential expression analysis of the normalized read counts between the sample groups was performed following best practices and recommendations of EdgeR^29,30^ and Limma^31^ on R environment (version 3.6.0). All the code used for data analysis is available upon request.

### Statistical analysis

No statistical method was used to predetermine sample size. The experiments were not randomized. The investigators were not blinded to allocation during experiments and outcome assessment. None of the samples/animals were excluded from the analysis. Data analyzed by parametric tests are represented by the mean_±_standard error of mean (S.E.M.) of pooled experiments unless otherwise stated. Values of *n* represent the number of independent experiments performed or the number of individual mice or patient specimens. For each independent *in vitro* experiment, at least 3 technical replicates were used, and a minimum number of three experiments were performed to ensure adequate statistical power. In the *in vitro* experiments, normal distribution was assumed, and one sample *t*-test was applied for one-component comparisons with control and Student’s *t*-test for two-component comparisons. For *in vivo* experiments, a non-parametric Mann-Whitney *U*-test was used. Two-tailed statistical analysis was applied for experimental design without predicted result, and one-tail for validation or hypothesis-driven experiments. The confidence level used for all the statistical analyses was 0.95 (alpha value_=_0.05). *p-value: * p<0.05, ** p<0.01, *** p<0.001*.

## Results

### Identification of metastatic prostate cancer-associated genes with prognostic potential

To unveil novel drivers of aggressive PCa with prognostic potential, we undertook an unbiased bioinformatics strategy taking advantage from transcriptomics data present in public datasets that contain tumor progression and patient outcome information (see workflow in Fig. 1A and cohort information in Fig. S1). We evaluated prognostic potential (the biochemical risk association - risk of recurrence of patients stratified according to the quartiles 1 vs. 4 of expression for each of the 22.021 genes present in 4 datasets that contained patient follow up information) (Fig.1A and Fig. S1) and selected genes that showed a consistent risk association between the interrogated datasets (202 genes). To prioritize genes that were associated to disease progression and not broadly to pathogenesis, we selected genes that were altered in the course of metastasis (comparing primary tumors and metastases) but not during initiation (excluding genes consistently altered in primary tumors vs. normal tissue). We applied stringent criteria to identify the candidates that showed a more robust trend between the interrogated 5 datasets. From the initial 21.357 genes, we shortlisted 77 upregulated and 175 downregulated genes. When integrating differential expression in metastasis and prognostic potential, 5 upregulated and 4 downregulated genes complied with all the above criteria (individual data for these genes is presented in Figure 1B).

**Fig 1.**
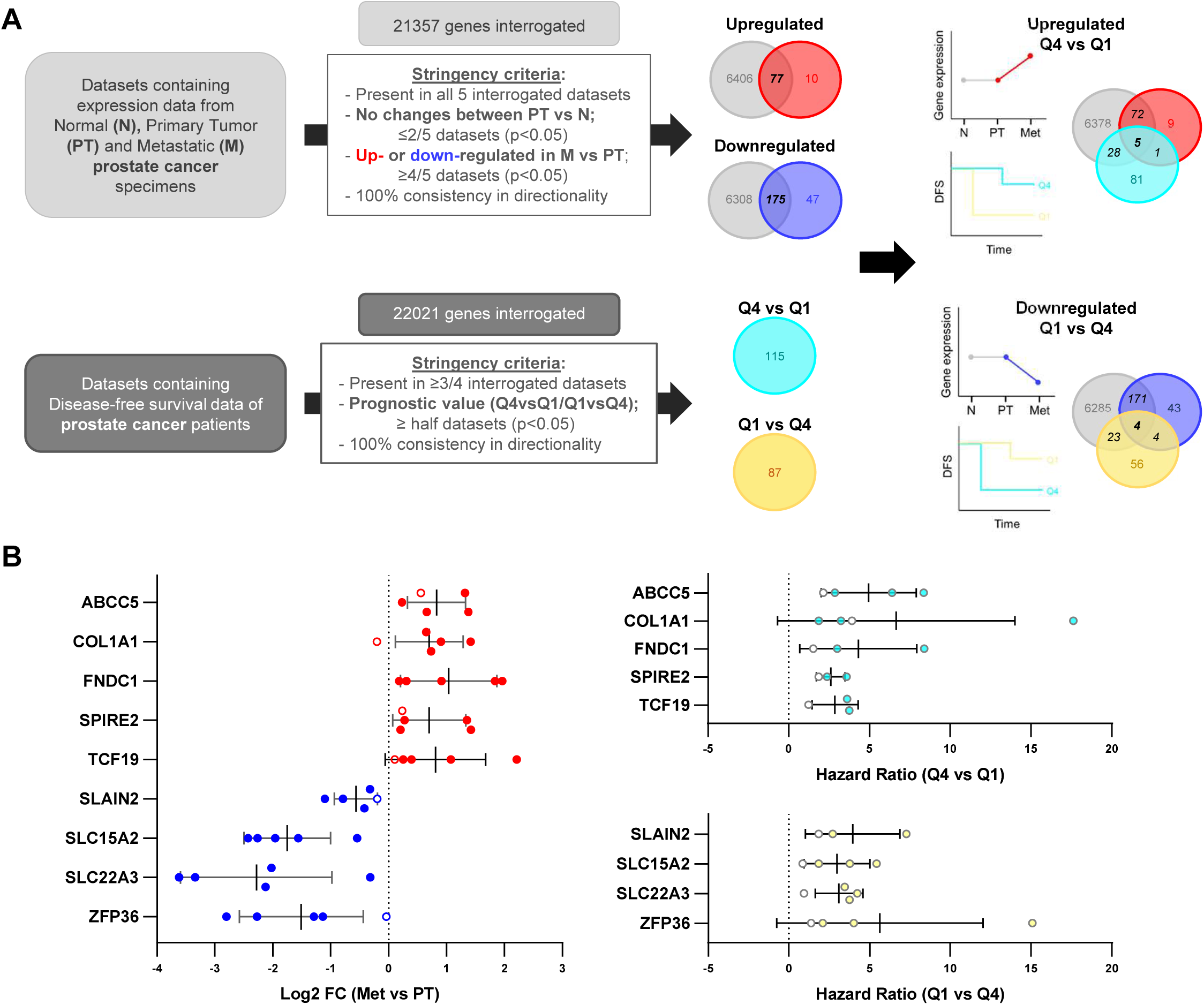
Identification of metastatic prostate cancer associated genes with prognostic potential. **A)** Workflow of the computational screening. The stringency criteria are indicated. The Log_2_-normalized gene expression was used to calculated the limma differential expression and select the up- or down-regulated genes. The prognostic value of the genes was analyzed by measuring the Hazard ratio (HR) between the Q1 and Q4 groups. Quartiles represent ranges of expression that divide the set of values into quarters being Q1 and Q4 the ones with higher and lower expression respectively. A Cox proportional hazards regression model was performed to compare the two groups. **B)** The Log_2_ Fold Change (FC) and Hazard ratio values of the candidate genes in each dataset are represented. The statistically significant results are represented with filled-colored dots.

### *TCF19* is a prognostic gene upregulated in metastatic prostate cancer patients

To select the gene best suited for molecular and biological studies, we focused on tumor cell intrinsic aggressiveness promoter genes (upregulated). We took advantage of single-cell portal to interrogate a single-cell RNA-seq dataset from PCa patients and identify the genes expressed in the epithelial compartment ^32^. Of the initial 9 genes, only *ABCC5* and *TCF19* met this criterion (Fig. 2A and Fig. S2). Of note, ABCC5 had been previously reported to contribute to PCa progression and androgen therapy resistance^33,34^, validating the potential of our screening strategy to uncover aggressive PCa associated genes (Fig. S3-S4). In contrast, the contribution of *TCF19* to PCa progression (which had a robust and consistent association to biochemical recurrence and metastasis, Fig. 2B, C) had never been addressed before, which prompted us to its study.

**Fig 2.**
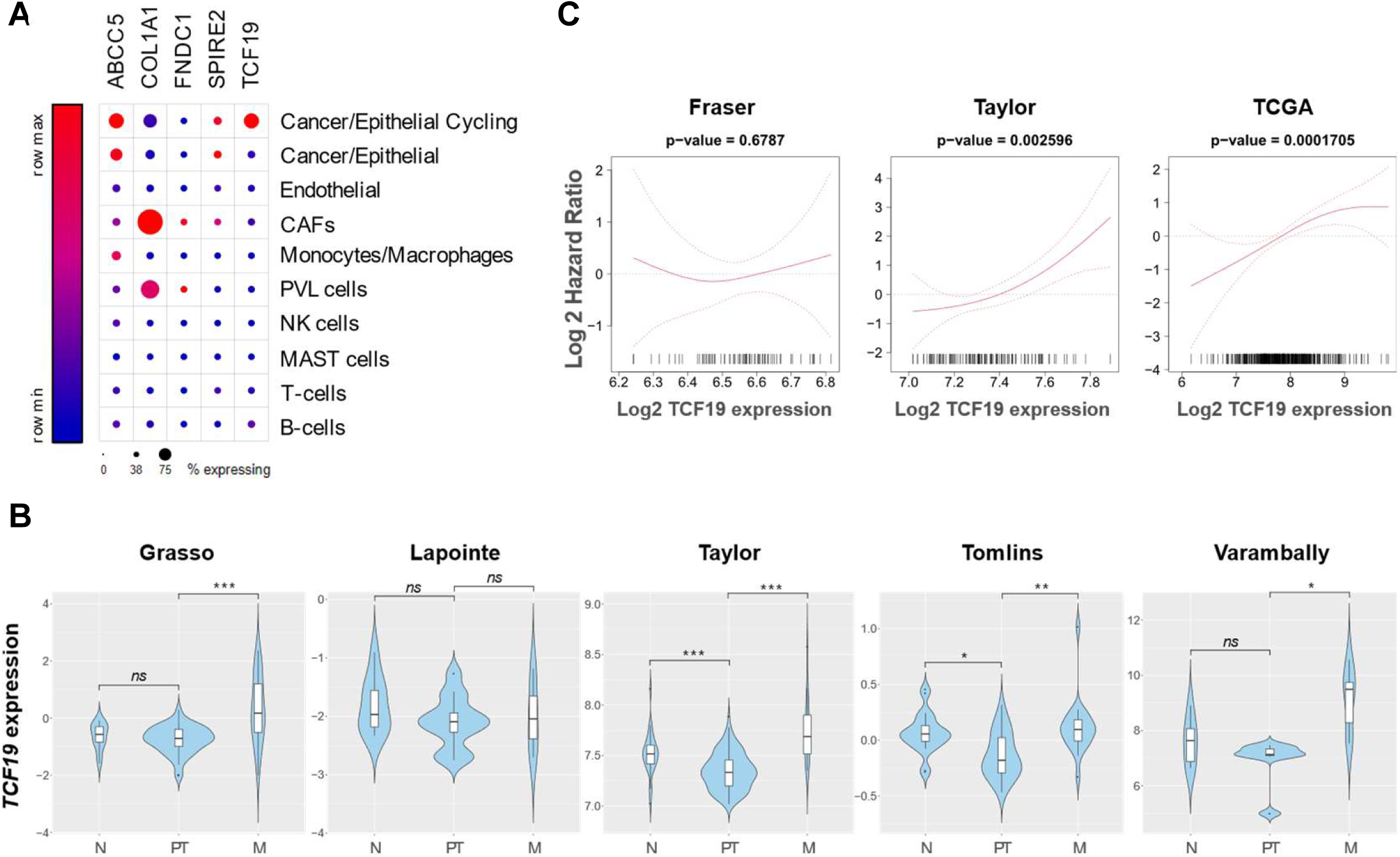
*TCF19* is a prognostic gene upregulated in metastatic prostate cancer patients. **A)** The relative expression of each gene in the indicated cell type was retrieved from the single cell data from a prostate cancer study (PMID: 33971952). The size of the dot represents the % of expressing cells. **B)** Violin plots depicting the expression of *TCF19* among non-tumoral (N), primary tumor (PT) and metastatic (M) prostate cancer specimens in the indicated datasets. The y-axis represents the Log_2_ -normalized gene expression (fluorescence intensity values for microarray data or, sequencing reads values obtained after gene quantification with RSEM and normalization using Upper Quartile in case of RNA-seq). *p* value derives from the limma differential expression between the indicated groups in the computational analysis *(p, p-value. ns p≥0.05, * p<0.05, ** p<0.005, *** p<0.0005).* **C)** Smooth Hazard Ratio curves. x-axis represents the gene expression level and y-axis the Log Hazard ratio. The p-value indicates the significance of the association between the gene and the outcome calculated via a likelihood ratio test.

### *TCF19* and androgen receptor target genes are inversely co-expressed in prostate cancer

*TCF19* is a growth regulated gene that has been increasingly associated with the proliferation potential of multiple cancer types^13–20^. Interestingly, our screening pointed at an unprecedented role for TCF19 in PCa dissemination. To identify pathways associated with the upregulation of TCF19 in PCa, we performed Gene Set Enrichment Analysis (GSEA)^23^. We stratified the patients of the 2 of the datasets that contained the higher number of individuals (Taylor (n=131) and TCGA (n=497)) in high or low TCF19 based on the mean *TCF19* expression (Fig. S5A). The GSEA analysis identified androgen response as the top repressed pathway in patients with elevated TCF19 mRNA expression (Fig. 3A-B). Of note, oxidative phosphorylation was also one of the top pathways consistent with previous reports^12^ and validating the strength of our approach. To confirm an inverse relationship between *TCF19* expression and AR signaling, we performed a correlation analysis between *TCF19* and an AR signature that is based on 31 genes previously developed by our group^35^. Of note, *TCF19* and AR signature showed a robust inverse correlation in Taylor and TCGA, and a similar trend in additional 2 datasets that contained a lower number of patients (Fig. 3C). Moreover, similar results were obtained by analyzing the correlation of *TCF19* with the AR target *KLK3* (Fig. S5B) and an additional public AR signature (Fig. S5C)^36^.

**Fig 3.**
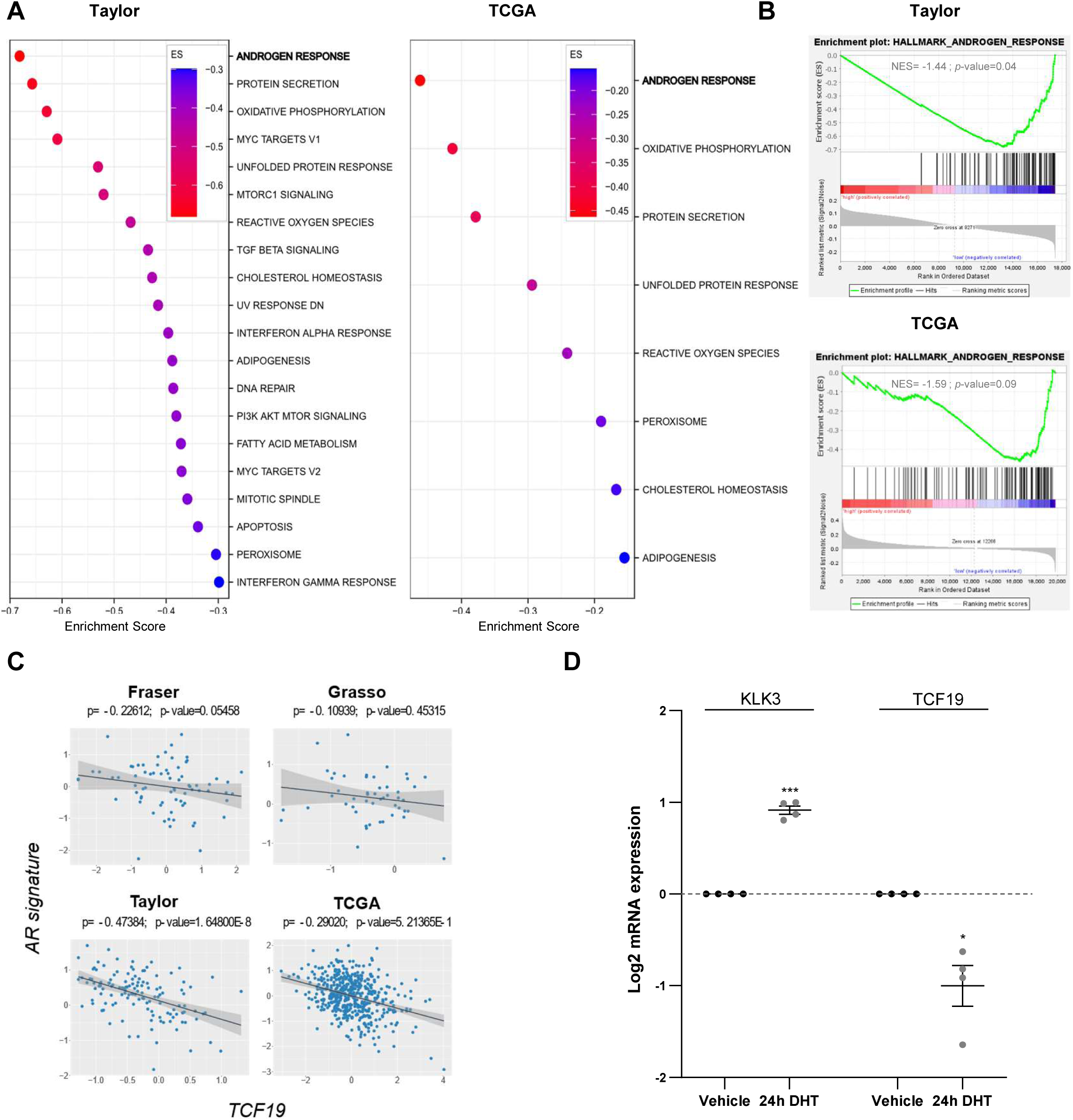
*TCF19* and androgen receptor target genes are inversely co-expressed in prostate cancer. **A and B)** Gene set enrichment analysis (GSEA) of TCF19 high vs.TCF19 low patients from Taylor and TCGA datasets. Primary tumor cohorts were divided in two groups based on their mean *TCF19* mRNA levels. The pathways associated with the levels of TCF19 and the Enrichments score (ES) plots of the most inversely correlated ones in each dataset are indicated. Cohort size is indicated in Fig S5A. **C)** Plotted values correspond to the Log_2_ - normalized gene expression values of *TCF19* and the androgen receptor (AR) signatures from (PMID: 34503116) (in X and Y-axis) in the primary tumor specimens from each patient in the indicated dataset. Black line represents linear regression, grey area indicates the limits of the confidence intervals and p and p-value indicate Spearman’s correlation coefficient and statistical significance respectively. **D)** Analysis of *KLK3* and *TCF19* expression by qRT-PCR in the AR-dependent LNCaP cells treated for 24h with AR agonist (dihydrotestosterone, DHT, 10 nM). Data were normalized to *GAPDH* expression and untreated (vehicle) condition. The dotted line represents the normalized value of the vehicle data. A one sample *t-*test was performed. *n* = 4 independent experiments. *p, p-value. ns p≥0.05, * p<0.05, ** p<0.01, *** p<0.001*.

AR has been commonly considered a transcriptional activator. Yet, increasing evidence supports a role of AR as a transcriptional repressor^3^. To assess whether *TCF19* could be repressed by AR, we treated AR proficient human prostate cancer cell lines with the AR agonist dihydrotestosterone (DHT)^37^. Interestingly, DHT-induced AR activation (Fig. 3D) resulted in the downregulation of *TCF19* in multiple PCa cell lines (Fig. 3D and Fig. S5D), suggesting that, directly or indirectly, AR represses *TCF19*. These results support the notion that primary tumors with lower androgen activity or tumors that evade the action of androgen deprivation therapy (ADT) and androgen receptor pathway inhibitors (ARPI)s would exhibit higher TCf19 expression levels.

### Context-dependent growth inhibition upon *TCF19* silencing in prostate cancer cells

TCF19 depletion has been reported to compromise the proliferation and foci formation capacity of colorectal^18^, head and neck squamous carcinoma^38^ and breast cancer^21^ cell lines. To assess whether *TCF19* silencing compromises also the proliferation potential of prostate cancer cell lines, we generated stable shRNA expressing PC3, DU145 and 22RV1 cells. The 2 different shRNAs used for these assays showed a similar silencing efficacy (Fig. 4A). Surprisingly, TCF19 silencing had a negligible effect in two-dimensional cell growth and cell cycle progression under unrestricted conditions (Fig. S6A, B). However, the growth of TCF19-silenced PC3, DU145 and 22RV1 cells in clonal conditions (colony formation assay) showed a consistent decrease in the foci forming ability (Fig. 4B). Moreover, TCF19-depleted cells also showed a decreased ability to grow in an anchorage independent manner in soft agar colony formation assays (Fig. 4C).

**Fig 4.**
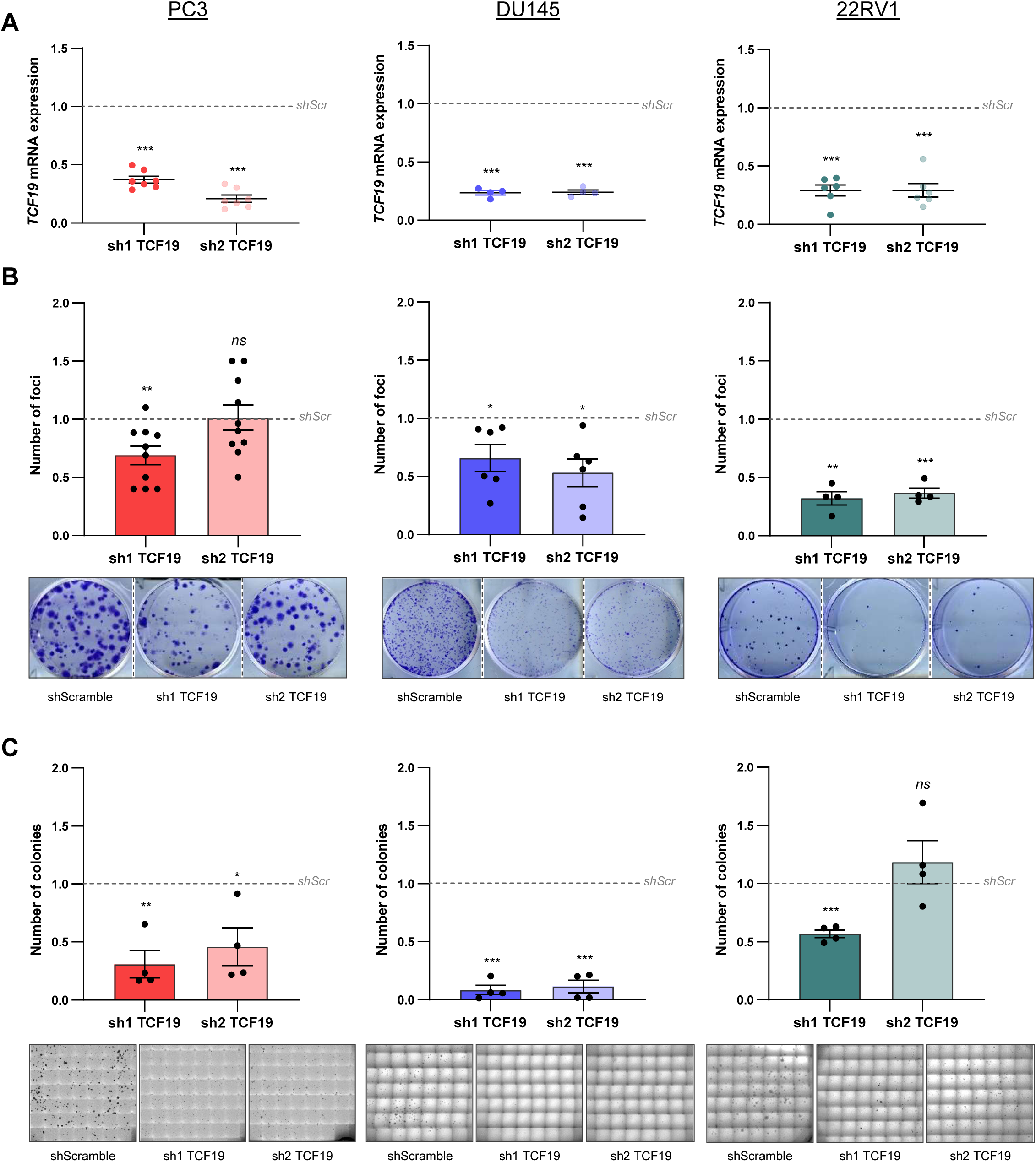
Context-dependent growth inhibition upon *TCF19* silencing in prostate cancer cells. **A)** Analysis of *TCF19* gene expression by qRT-PCR in the indicated cell line upon silencing of *TCF19* by shRNA transduction. Data were normalized to *GAPDH* expression and shScramble (shScr) condition. The dotted line represents the normalized value of the shScramble data. A one sample *t-*test was performed for statistical analysis. PC3 (*n*=7), DU145 (*n*=4) and 22RV1 (*n*=6) independent experiments **B)** Analysis of foci formation upon *TCF19* depletion. The number of foci normalized to shScramble condition are shown (top panels). A one sample *t*-test was performed for statistical analysis. PC3 (*n*=10), DU145 (*n*=6) and 22RV1 (*n*=4) independent experiments. Representative images are shown (bottom panels). **C)** Analysis of anchorage independent growth. The number of colonies normalized to shScramble are shown (top panels). A one sample *t*-test was performed for statistical analysis. (*n*=4, independent experiments). Representative images are shown (bottom panels). *p, p-value. ns p≥0.05, * p<0.05, ** p<0.01, *** p<0.001*.

Alterations in the migration and invasion ability of TCF19-depleted cells have also been reported^18,21,38^. However, we did not observe consistent defects on the migration and invasion ability of human PC3 prostate cancer cells (Fig. S6C-D).

### TCF19 depletion compromises tumor growth and metastatic capacity of prostate cancer cells

In contrast to previous reports, the effect of *TCF19* silencing in human prostate cancer cell lines did not show a profound impact on 2D assays (Fig. 4 and Fig. S6). This fact, together with the association of *TCF19* to PCa dissemination emerging from our bioinformatics screening led us to hypothesize that the role of TCF19 in PCa might go beyond promoting cellular proliferation. Recent reports have highlighted tumorigenic roles of TCF19 in pathways different from cellular proliferation^21,22^. To address whether this was also the case in the context of PCa, we performed two complementary *in vivo* studies. On the one hand, we performed an *in vivo* orthotopic assay (Fig. 5A). To this end, we generated stable shRNA expressing GFP-Luc PC3 cells and injected them into the ventral lobe of the prostates of immune-deficient nude mice. Local tumor growth and metastatic outgrowth were monitored following the Luciferase expressing cells by IVIS. Remarkably, TCF19-depleted cells showed a strong defect in tumor growth that correlated with a reduced primary tumor size and weight at the experimental endpoint (Fig. 5B and Fig. S7A). Moreover, the analysis of luciferase signal in distal organs *ex vivo* confirmed a lower metastatic burden in the bones of mice injected with *TCF19*-silenced PCa cells (Fig. 5C and Fig. S7B). On the other hand, we performed a complementary *in vivo* metastatic assay. For this second approach, we transduced again GFP-Luc expressing PC3 cells with TCF19-targeting or Scramble shRNA and injected them intracardially in the left ventricle of immune-deficient nude mice and monitored the luciferase signal over 27 days (Fig. 5D). Remarkably, *TCF19*-silenced cells showed lower number of bone lesions (Fig. 5E), confirming a role of TCF19 in sustaining metastatic dissemination in PCa.

**Fig 5.**
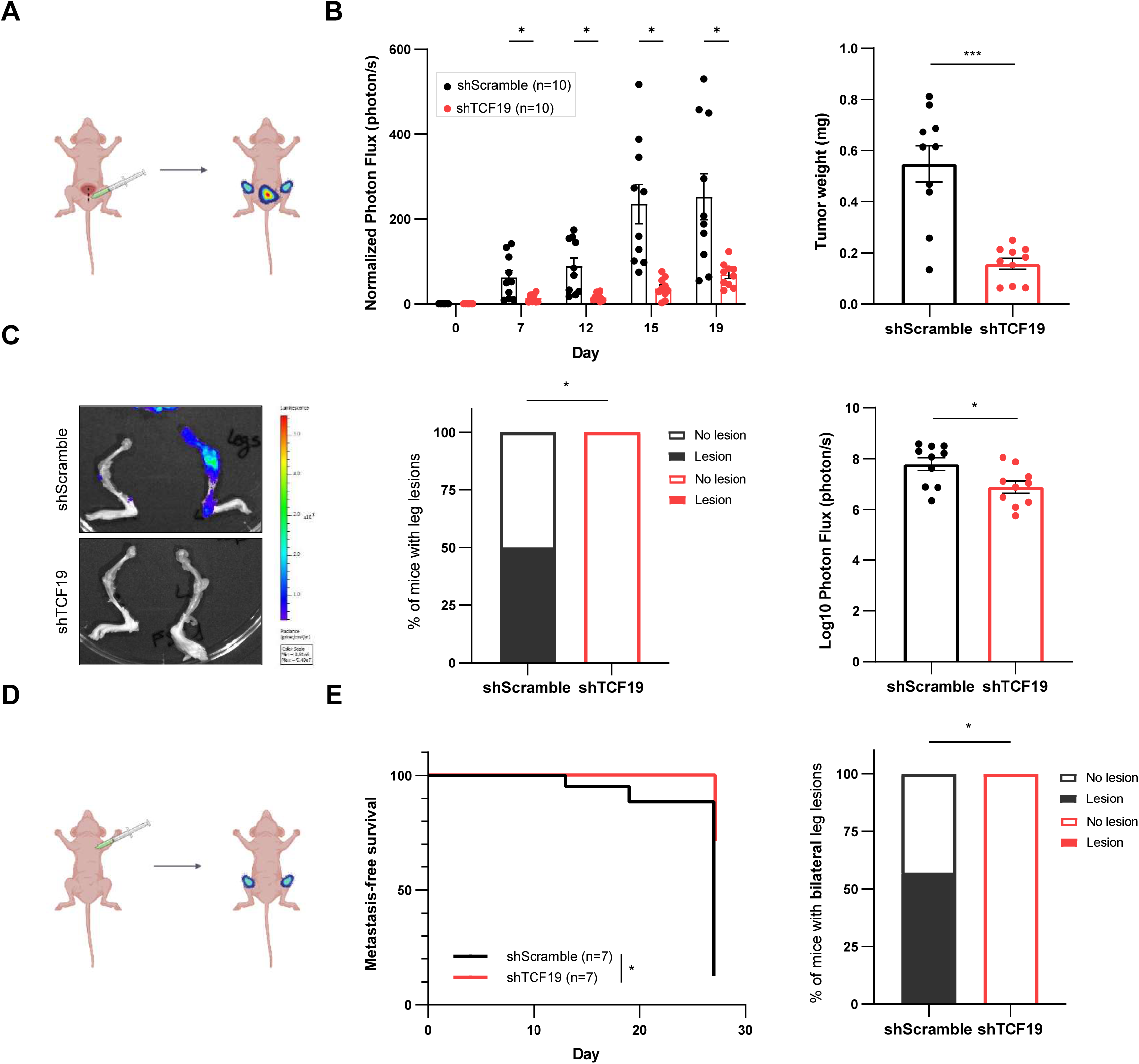
TCF19 depletion compromises tumor growth and metastatic capacity of prostate cancer cells. **A)** Schematic representation of the orthotopic xenotransplant assay performed to assess the local tumor growth and the metastatic capacity of *TCF19*-depleted PC3 GFP-Luc cells (*n*=10 mice/group). **B)** Evaluation of local tumor growth by orthotopic xenotransplant assay. IVIS relative flux data along the experimental process (left panel). The total photon flux normalized to time 0 are represented. A multiple Mann-Whitney *U*-test was applied for statistical analysis. *Ex vivo* tumor weight of the ventral lobes of the prostates are represented (right panel). A one-tailed unpaired Student’s *t*-test was applied for statistical analysis. **C)** Evaluation of metastatic lesions in the legs by orthotopic xenotransplant assay. Representative images (left panel). The *ex vivo* incidence of leg lesions are represented (middle panel). Luciferase signal above day 0 was considered metastasis-positive. The Log_10_ photon flux of the sum signals from both legs are represented (right panel). A two-sided Fisher’s exact test was performed. **D)** Schematic representation of the intra-cardiac xenotransplant assay performed to assess the metastatic capacity of *TCF19*-depleted PC3 GFP-Luc cells (*n*=7 mice/group). **E)** Metastasis-free survival curves of prostate cancer cell signal in legs was monitored for up to 27 days (left panel). Luciferase signal above day 0 was considered metastasis-positive. The incidence of bilateral leg lesions at day 27 are represented (right panel). A log-rank test and a one-sided Fisher’s exact test were performed respectively. *p, p-value. ns p≥0.05, * p<0.05, ** p<0.01, *** p<0.001*.

### *TCF19* silencing in prostate tumors results in a decrease in hypoxia-responsive genes

To address the molecular basis of the activity of TCF19 in PCa, we first focused on the reported association with FOXO1 regulation^14,16^. To assess whether alterations in FOXO transcriptional program were present in TCF19-depleted tumors (Fig. 5) we analyzed the mRNA levels of representative targets of this transcription factor in specimens collected from the *in vivo* orthotopic assay. *p27* and *CCND1* mRNA levels were unperturbed, and only *p21* gene expression was elevated in TCF19-silenced tumors, ruling out a major perturbation in FOXO activity (Fig. S8A). Consistently, TCF19 depletion did not cause a significant change in the proliferation markers Ki67 or pRb^pS807/811^ *in vivo* analyzed by immunofluorescence (Fig. S8B). The analysis of *VIMENTIN* and *SNAIL* mRNA levels in those tumors also ruled out major alterations in EMT (Fig. S8C) that contribute to metastasis in breast cancer^21^. We did not observe an increase in the apoptosis marker Cleaved-Caspase 3 either in TCF19-depleted tumors (Fig. S8D).

The analysis of previously reported processes did not shed light into the role of TCF19 in sustaining PCa metastasis. Hence, we decided to perform a high-throughput transcriptomics analysis from our tumor specimens, taking advantage of the primary tumor samples from the orthotopic assay. The heatmap of the top 200 differentially expressed genes (DEG) nicely grouped the tumors from each experimental condition (Fig. S9A). By applying an FDR<0.05 and a Log_2_FC|1| as cutoff, we identified 599 DEGs (GSE287409, Table S3). Of note, *TCF19* appeared downregulated in our analysis, which served as quality control (Fig. 6A). To identify the main pathways altered upon depletion of TCF19, we performed a GSEA analysis. Our GSEA analysis confirmed alterations in known TCF19-related molecular programs such as interferon signaling^22^, OXPHOS and glycolysis^12^ (Fig. 6B and Fig. S9B), but also identified novel potential pathways that could explain the defects observed in metastasis. Hypoxia responsive gene signatures are associated to biochemical recurrence^39^ and hypoxia promotes the acquisition of castration-resistant features in the context of PCa^40^. Interestingly, our GSEA analysis unveiled hypoxia as one of the main pathways affected by TCF19 depletion (Fig. 6B and Fig. S9C). We confirmed this data through the analysis of hypoxia target gene expression (*CA9* and *VEGFA*) in TCF19-depleted tumors (Fig. 6C). A similar trend was observed in three additional hypoxia target genes (*ANGPTL4, BNIP3, SLC2A1*; Fig. S9D). Moreover, the computational dissection of the tumor microenvironment in the *in vivo* orthotopic tumors using xCell^24^ uncovered endothelial cells as the stromal components exhibiting the most robust direct association with TCF19 perturbation (Fig 6D).

**Fig 6.**
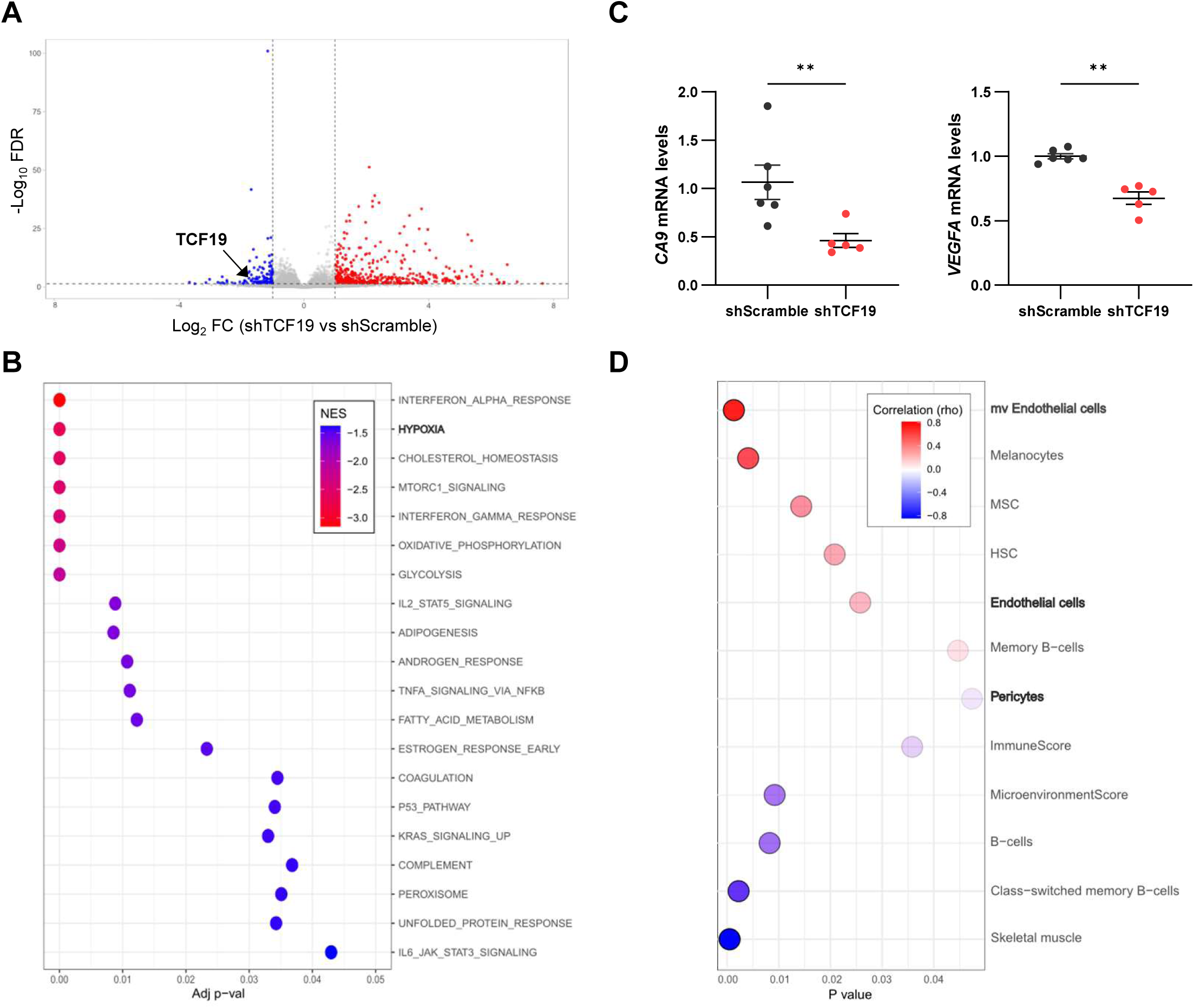
*TCF19* silencing in prostate tumors results in a decrease in hypoxia-responsive genes. **A)** Volcano plot representation of the differentially expressed genes (FDR<0.05; Log_2_FC>I1I) in the tumors collected from the orthotopic xenotransplant assay. The dot representing *TCF19* expression is highlighted. **B)** A GSEA analysis was performed to identify pathways enriched upon *TCF19* depletion. The Top 20 pathways phenotypically enriched in the control at the GSEA analysis are shown. **C)** Gene expression analysis of shScramble and shTCF19 tumors from the *in vivo* orthotopic experiment. The tumors selected for the qRT-PCR analysis are indicated in Fig S7. Data were normalized to *RPLP0* expression and shScramble condition. A Mann-Whitney *U*-test was performed for statistical analysis. **D)** Digital dissection of the tumor microenvironment in the *in vivo* orthotopic tumors from Fig. 5 using xCell (PMID: 29141660). The stromal compartments positively and negatively correlated with TCF19 were inferred from the RNA-seq data. *p, p-value. ns p≥0.05, * p<0.05, ** p<0.01, *** p<0.001*.

### *TCF19*-silenced tumors show vessel normalization with reduced vascular permeability

To explore whether the hypoxia and endothelial cell processes identified in molecular analyses translated to vascular alterations, we analyzed the tumors collected from the orthotopic xenotransplant assay (Fig. S7) by immunofluorescence. Endothelial abundance in the tumor area was estimated by means of CD31 staining. Counterintuitively, TCF19-depleted tumors showed an increased number of blood vessels compared to control ones (Fig. 7A and Fig. S10A). However, they exhibited distinctive structural and functional features indicative of increased functionality. We monitored the pericyte compartment in vessels to address their maturity. Interestingly, TCF19-depleted tumors showed an increased level of αSMA around the vessels, suggestive of increased functional pericyte coverage and a potential reduction in vascular permeability that enables tumor cell intravasation^41^. In turn, we analyzed vessel permeability indicators. On the one hand, we monitored endothelial junction tightening using the marker B-catenin, which was elevated suggesting tighter endothelial cell junctions (Fig. 7B). On the other hand, we analyzed erythrocyte extravasation in tumoral areas, which would be indicative of elevated vascular permeability, and found a profound reduction in extravascular Ter119 staining (Fig. 7C). Altogether the characterization of the vascular bed in tumors revealed a regulation of vascular permeability and function upon TCF19 depletion, which is consistent with the reduced growth and dissemination of PCa.

**Fig 7.**
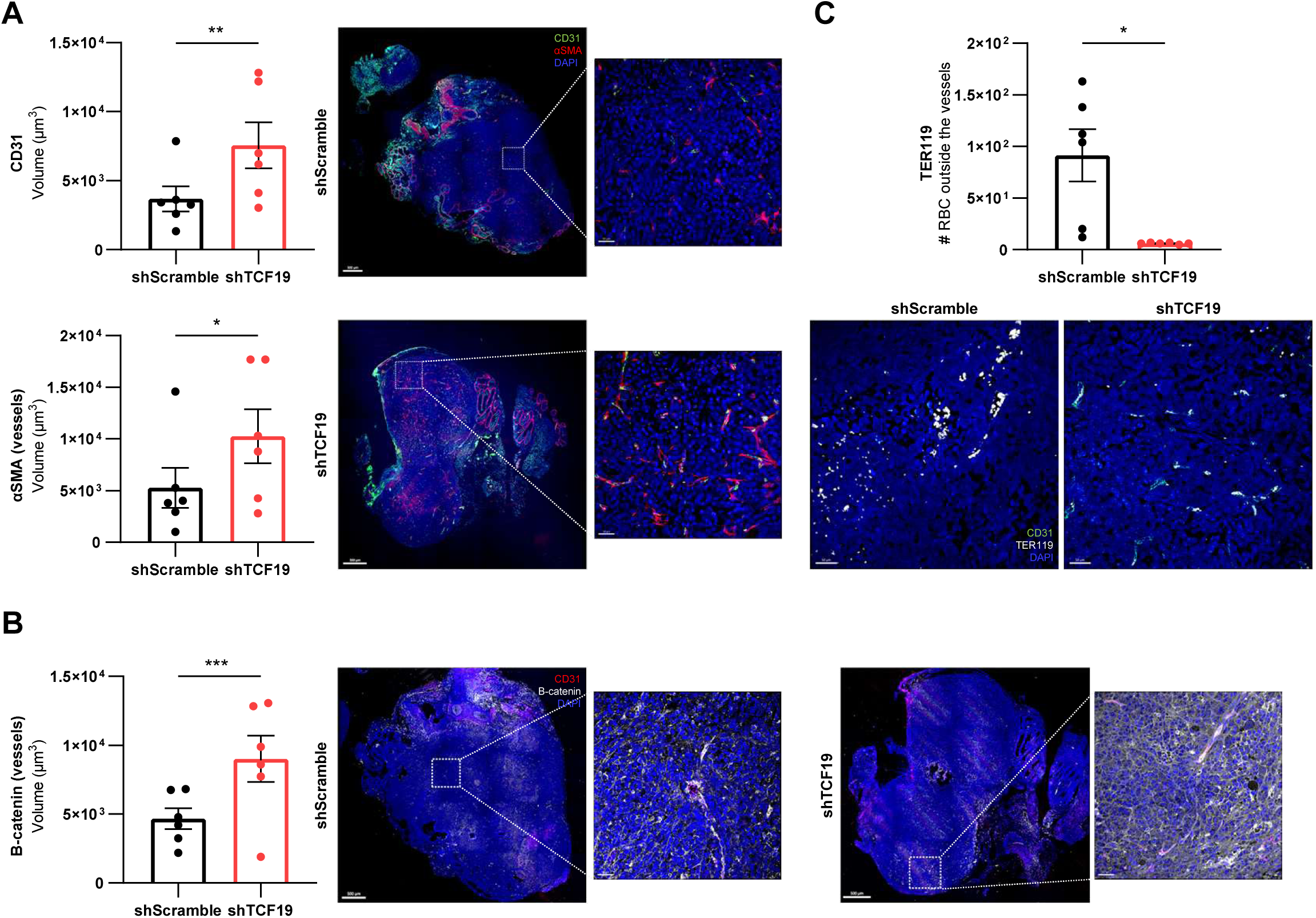
TCF19-silenced tumors show vessel normalization with reduced vascular permeability. Analysis of the indicated proteins by immunofluorescence in shScramble and shTCF19 tumors from the *in vivo* orthotopic experiment. The tumors selected for the IHC/IF analysis are indicated in Fig S7. A Nested *t*-test was performed for statistical analysis (left panels). Representative images are shown (right panels)*. p, p-value. ns p≥0.05, * p<0.05, ** p<0.01, *** p<0.001*.

## Discussion

Recent advances in PCa research have increased our understanding of this complex disease that still causes more than 350.000 annual deaths, which are mainly due to the acquisition of androgen resistant and chemorefractory features^1^. Uncovering the key determinants of PCa aggressiveness is essential to understand the high morbidity due to metastatic disease.

By analyzing patient data from public repositories, we have identified 9 genes with prognostic potential that are dysregulated in metastatic PCa patients. The genes identified in our screening are expressed in epithelial and stromal cells, suggesting that both tumor cell-intrinsic and extrinsic alterations can be indicative and/or contribute to the gain of aggressive and metastatic features.

*TCF19* ranked as a top upregulated epithelial gene in our screening. The contribution of TCF19 to tumor progression by controlling cellular proliferation has been reported in gastric cancer^13^, hepatocellular carcinoma^14,15^, non-small cell lung cancer^16,17^, colorectal cancer^18^ and liver cancer^19^. In addition, TCF19 levels have been associated with the risk of suffering from head and neck cancer^38^ and it has proven prognostic potential for colorectal cancer^18^, renal clear cell carcinoma^42^, endometrial cancer^22^ and hepatocellular carcinoma^43^ patients. Consistent with this, our patient data shows a tumor and metastasis supportive role for TCF19 in PCa. Interestingly, the combination of computational, *in vitro* and *in vivo* experimental strategies revealed that beyond the role of TCF19 in supporting tumor-intrinsic properties, it also contributes to tumor microenvironment remodeling. Although the exact molecular mechanism driving this phenomenon remain to be elucidated, our work provides evidence for the role of tumor cell-expressed *TCF19* in vessel homeostasis and permeability, a process that is critical in cancer cell dissemination^44^.

Inhibition of androgen signaling or AR activity elicits a transient clinical benefit, that is bypassed in patients leading to CRPC. Importantly, the mechanisms that enable prostate cancer cell adaptation to lack of AR signaling are poorly understood but could include transcriptional rewiring and cellular plasticity^1^. Indeed, a gene that ranked as a top prognostic factor in PCa progression in our screening, ABCC5, is reportedly associated to the acquisition of CRPC features^33,34^. Our results demonstrate that AR signaling opposes *TCF19* expression. In turn, it is tempting to speculate that the upregulation of TCF19 upon treatment with ADT or AR inhibitors could represent a first adaptive response that would support prostate cancer cell function and the eventual acquisition of aggressive features. In line with our findings of TCF19-regulated hypoxia response and vessel function, the reported contribution of hypoxia to biochemical recurrence^39^ and to the acquisition of castration-resistant features^40^ could be indicative of an important role for the TCF19-hypoxia-vessel homeostasis in PCa progression. Future work will clarify the real contribution of TCF19 to the gain of castration-resistant features.

### Conclusions

In this study, we combine computational and empiric approaches to identify the causal role and prognostic potential of *TCF19* in aggressive prostate cancer. We find that this gene is repressed by androgen signaling and contributes to stroma remodeling by means of vessel homeostasis and permeability, a central process in cancer cell dissemination.

## Acknowledgements

We are grateful to the members of the Torrano and the Carracedo labs for discussions and technical advice. We thank Kathrin Keim for administrative support and valuable input. We are grateful to Monika Gonzalez-Lopez and José Ezequiel Martín from the Genome Analysis Platform at CIC bioGUNE for technical support.

## Author contributions: CRediT

AE designed and performed the majority of experiments, data analysis, prepared the figures and drafted the manuscript. JR-C, MF and S-FR provided technical support with the *in vitro* experiments. SG-L and IM provided bioinformatics support. NM-M contributed to the design and data analysis of computational screening. OC, IA, and MGu provided technical support with *in vivo* experiments. The RNA-seq analysis was performed at the Genome Analysis Platform at CIC bioGUNE with the support of LB and AM-A. MGr and RR-G contributed to experimental design and provided valuable input. AC conceived the study, supervised the execution of the project and drafted the manuscript. All authors have read and approved the final version of the manuscript.

## Ethics Statement

The authors declare no competing interest.

## Data Availability Statement

The authors declare that data supporting the findings of this study are available within the paper and its supplementary files. RNA-seq data is accessible in GEO repository with ID GSE287409.

## References

1. Rebello, R. J., et al. Prostate cancer. Nat Rev Dis Primers 7, (2021).

2. Martínez-Jiménez, F. et al. Pan-cancer whole-genome comparison of primary and metastatic solid tumours. Nature 2023 618:7964 618, 333–341 (2023).

3. Chen1, R., et al. Transcriptional repression by androgen receptor: roles in castration-resistant prostate cancer. Asian J Androl 21, 1–4 (2019).

4. Ku, D. H. et al. A new growth-regulated complementary DNA with the sequence of a putative trans-activating factor. Cell growth & differentiation 2, 179–186 (1991).

5. Sen, S. et al. Transcription factor 19 interacts with histone 3 lysine 4 trimethylation and controls gluconeogenesis via the nucleosome-remodeling-deacetylase complex. Journal of Biological Chemistry 292, 20362–20378 (2017).

6. Cheung, Y. H., Watkinson, J. & Anastassiou, D. Conditional meta-analysis stratifying on detailed HLA genotypes identifies a novel type 1 diabetes locus around TCF19 in the MHC. Hum Genet 129, 161–176 (2011).

7. Ndiaye, F. K. et al. Expression and functional assessment of candidate type 2 diabetes susceptibility genes identify four new genes contributing to human insulin secretion. Mol Metab 6, 459–470 (2017).

8. Harder, M. N. et al. The type 2 diabetes risk allele of TMEM154-rs6813195 associates with decreased beta cell function in a study of 6,486 danes. PLoS One 10, 1–13 (2015).

9. Barbosa-Sampaio, H. C. et al. Nupr1 deletion protects against glucose intolerance by increasing beta cell mass. Diabetologia 56, 2477–2486 (2013).

10. Krautkramer, K. A. et al. Tcf19 is a novel islet factor necessary for proliferation and survival in the INS-1 β-cell line. Am J Physiol Endocrinol Metab 305, 600–610 (2013).

11. Yang, G. H. et al. Tcf19 impacts a network of inflammatory and dna damage response genes in the pancreatic β-cell. Metabolites 11, (2021).

12. Mondal, P. et al. TCF19 and p53 regulate transcription of TIGAR and SCO2 in HCC for mitochondrial energy metabolism and stress adaptation. The FASEB Journal 35, 1–23 (2021).

13. Miao, R. et al. VEZT, a Novel Putative Tumor Suppressor, Suppresses the Growth and Tumorigenicity of Gastric Cancer. PLoS One 8, 1–14 (2013).

14. C.X. zeng, et al. TCF19 enhances cell proliferation in hepatocellular carcinoma by activating the ATK/FOXO1 signaling pathway. Neoplasma 26, 300–303 (2018).

15. Mondal, P. et al. TCF19 promotes cell proliferation through binding to the histone H3K4me3 mark. Biochemistry (2019) doi:10.1021/acs.biochem.9b00771.

16. Zhou, Z. H. et al. TCF19 contributes to cell proliferation of non-small cell lung cancer by inhibiting FOXO1. Cell Biol Int 43, 1416–1424 (2019).

17. Tian, Y. et al. TCF19 promotes cell proliferation and tumor formation in lung cancer by activating the Raf/MEK/ERK signaling pathway. Transl Oncol 45, (2024).

18. Du, W. et al. TCF19 aggravates the malignant progression of colorectal cancer by negatively regulating WWC1. ERMPS 655–663 (2020).

19. Xu, G., Zhu, Y., Liu, H., Liu, Y. & Zhang, X. Lncrna mir194-2hg promotes cell proliferation and metastasis via regulation of mir-1207-5p/ tcf19/wnt/β-catenin signaling in liver cancer. Onco Targets Ther 13, 9887–9899 (2020).

20. Tan, J., lian, H., Zheng, Q., Yang, T. & Wang, T. TCF19 Enhances Glioma Cell Proliferation via Modulating the Β-Catenin Signaling Pathway through Accelerating DHX32 Transcription. Curr Cancer Drug Targets 25, (2025).

21. Li, S. yu, et al. Deciphering the TCF19/miR-199a-5p/SP1/LOXL2 pathway: Implications for breast cancer metastasis and epithelial-mesenchymal transition. Cancer Lett 597, (2024).

22. Ma, X. et al. Targeting TCF19 sensitizes MSI endometrial cancer to anti-PD-1 therapy by alleviating CD8+ T cell exhaustion via TRIM14-IFN-β axis. Cell Rep 42, (2023).

23. Subramanian, A. et al. Gene set enrichment analysis: a knowledge-based approach for interpreting genome-wide expression profiles. Proc Natl Acad Sci U S A 102, 15545–15550 (2005).

24. Aran, D., Hu, Z. & Butte, A. J. xCell: digitally portraying the tissue cellular heterogeneity landscape. Genome Biol 18, (2017).

25. Crespo, J. R. et al. The PP2A regulator IER5L supports prostate cancer progression. Cell Death Dis 15, 514 (2024).

26. Dobin, A. et al. STAR: ultrafast universal RNA-seq aligner. Bioinformatics 29, 15–21 (2013).

27. Danecek, P. et al. Twelve years of SAMtools and BCFtools. Gigascience 10, (2021).

28. Liao, Y., Smyth, G. K. & Shi, W. featureCounts: an efficient general purpose program for assigning sequence reads to genomic features. Bioinformatics 30, 923–930 (2014).

29. Chen, Y., Lun, A. T. L. & Smyth, G. K. From reads to genes to pathways: Differential expression analysis of RNA-Seq experiments using Rsubread and the edgeR quasi-likelihood pipeline. F1000Res 5, (2016).

30. Robinson, M. D., McCarthy, D. J. & Smyth, G. K. edgeR: a Bioconductor package for differential expression analysis of digital gene expression data. Bioinformatics 26, 139–140 (2010).

31. Ritchie, M. E. et al. limma powers differential expression analyses for RNA-sequencing and microarray studies. Nucleic Acids Res 43, e47 (2015).

32. Wu, S. Z. et al. Cryopreservation of human cancers conserves tumour heterogeneity for single-cell multi-omics analysis. Genome Med 13, 1–17 (2021).

33. Ji, G. et al. Upregulation of ATP Binding Cassette Subfamily C Member 5 facilitates Prostate Cancer progression and Enzalutamide resistance via the CDK1-mediated AR Ser81 Phosphorylation Pathway. Int J Biol Sci 17, 1613–1628 (2021).

34. Chen, H. et al. Non-drug efflux function of ABCC5 promotes enzalutamide resistance in castration-resistant prostate cancer via upregulation of P65/AR-V7. Cell Death Discov 8, (2022).

35. Camacho, L. et al. Identification of Androgen Receptor Metabolic Correlome Reveals the Repression of Ceramide Kinase by Androgens. Cancers (Basel) (2021).

36. Hieronymus, H. et al. Gene expression signature-based chemical genomic prediction identifies a novel class of HSP90 pathway modulators. Cancer Cell 10, 321–330 (2006).

37. Tran, C. et al. Development of a second-generation antiandrogen for treatment of advanced prostate cancer. Science 324, 787–790 (2009).

38. Ji, P. et al. Genetic variants associated with expression of TCF19 contribute to the risk of head and neck cancer in Chinese population. J Med Genet (2021) doi:10.1136/jmedgenet-2020-107410.

39. Milosevic, M. et al. Tumor hypoxia predicts biochemical failure following radiotherapy for clinically localized prostate cancer. Clin Cancer Res 18, 2108–2114 (2012).

40. Cameron, S. et al. Chronic hypoxia favours adoption to a castration-resistant cell state in prostate cancer. Oncogene 42, 1693–1703 (2023).

41. Gerhardt, H. & Betsholtz, C. Endothelial-pericyte interactions in angiogenesis. Cell Tissue Res 314, 15–23 (2003).

42. Cheng, X. et al. Immunotherapeutic Value of Transcription Factor 19 (TCF19) Associated with Renal Clear Cell Carcinoma⍰: A Comprehensive Analysis of 33 Human Cancer Cases. J Oncol (2022).

43. Wang, M. et al. Transcription factors-related molecular subtypes and risk prognostic model: exploring the immunogenicity landscape and potential drug targets in hepatocellular carcinoma. Cancer Cell Int 24, 9 (2024).

44. Liu, Z. L., Chen, H. H., Zheng, L. L., Sun, L. P. & Shi, L. Angiogenic signaling pathways and anti-angiogenic therapy for cancer. Signal Transduct Target Ther 8, (2023).

