## Supplementary Figures and Figure legends for "Transcription factor 19 is an androgen responsive gene that modulates vessel homeostasis and sustains metastatic prostate cancer"

### Figure S1

| Dataset | Cohort size | | |
|---|---|---|---|
| | *Normal* | *Primary Tumor* | *Metastasis* |
| Grasso | 12 | 49 | 27 |
| Lapointe | 9 | 13 | 4 |
| Taylor | 29 | 131 | 19 |
| Tomlins | 23 | 32 | 20 |
| Varambally | 6 | 7 | 6 |

| Dataset | DFS.Status = 0 | DFS.Status = 1 | Missing data - with respect to PT | DFS.TIME min | DFS.TIME max | DFS.TIME mean |
|---|---|---|---|---|---|---|
| Fraser | 57 | 16 | 0 | 1.64 | 154.45 | 74.19 |
| Glinsky | 42 | 37 | 0 | 1.4 | 105.7 | 51.54 |
| Taylor | 104 | 27 | 0 | 1.38 | 149.19 | 48.19 |
| TCGA | 400 | 91 | 6 | 0.76 | 165.05 | 32.14 |

**Fig S1.** *Related to Fig 1*. Information related to the cohort size and Disease-Free Survival (DFS) of the datasets (PMID: 22722839) (PMID: 14711987) (PMID: 20579941) (PMID: 17173048) (PMID: 16286247) (PMID: 28068672) (PMID: 15067324) (https://gdac.broadinstitute.org/) used for the computational screening.

### Figure S2

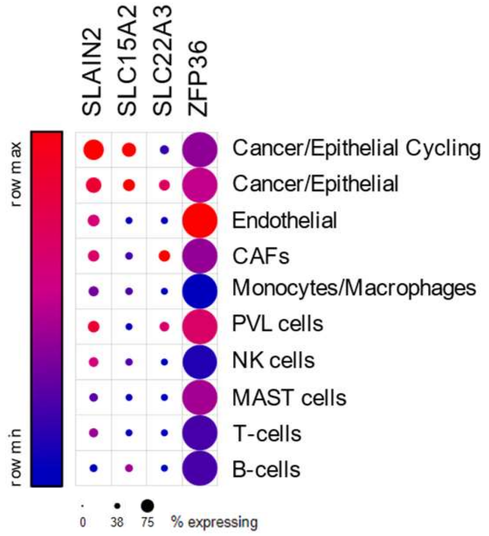


**<u>Fig S2.</u>** *Related to Fig 2*. The relative expression of each gene in the indicated cell type was retrieved from the single cell data from a prostate cancer study (PMID: 33971952). The size of the dot represents the % of expressing cells.

### Figure S3

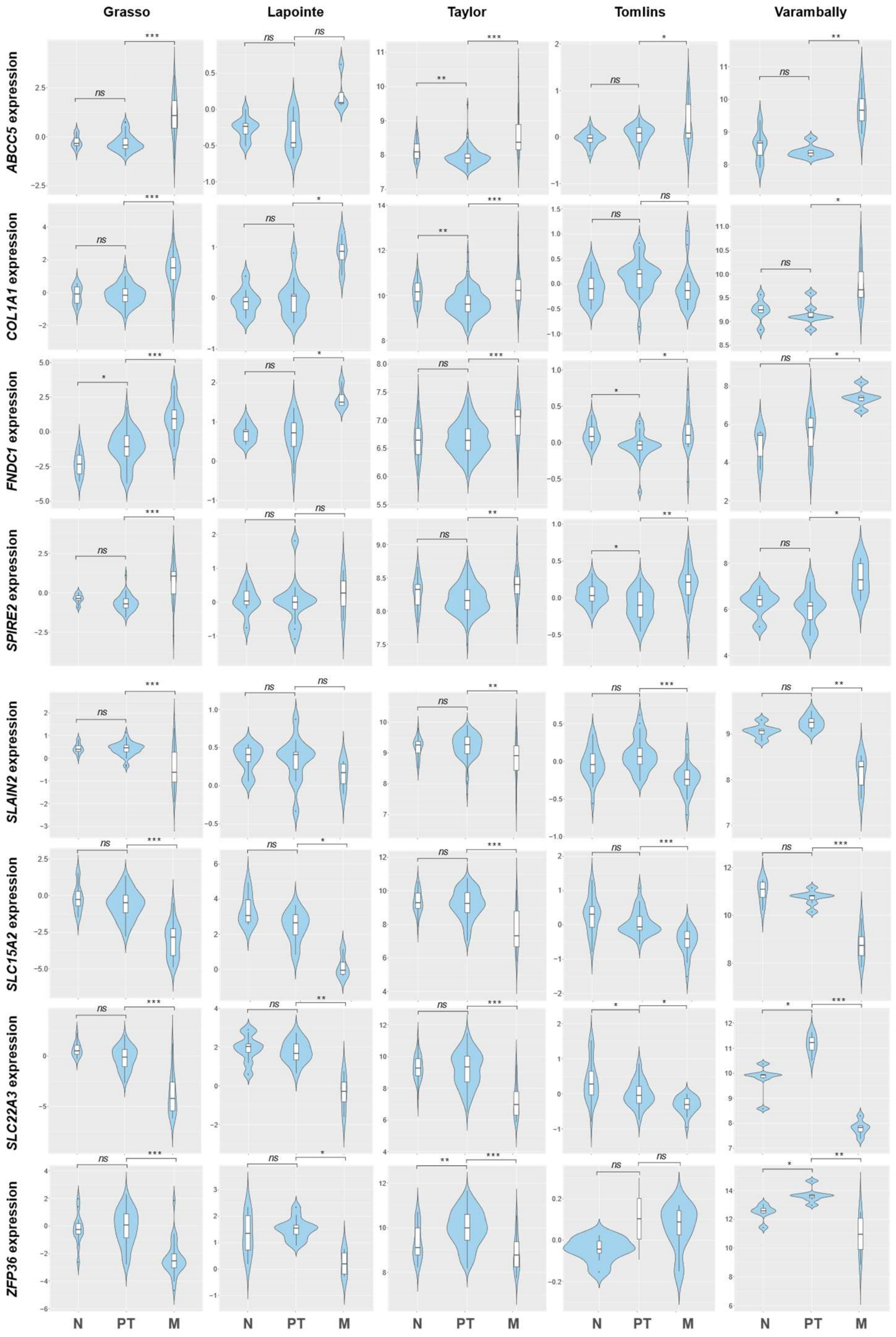


**Fig S3.** *Related to Fig 2*. Violin plots depicting the expression of the indicated gene among non-tumoral (N), primary tumor (PT) and metastatic (M) prostate cancer specimens in the indicated datasets. The y-axis represents the $Log_2$-normalized gene expression (fluorescence intensity values for microarray data or, sequencing reads values obtained after gene quantification with RSEM and normalization using Upper Quartile in case of RNA-seq). p value derives from the limma differential expression between the indicated groups in the computational analysis *(p,p-value. ns p≥0.05, * p<0.05, ** p<0.005, *** p<0.0005).*

### Figure S4

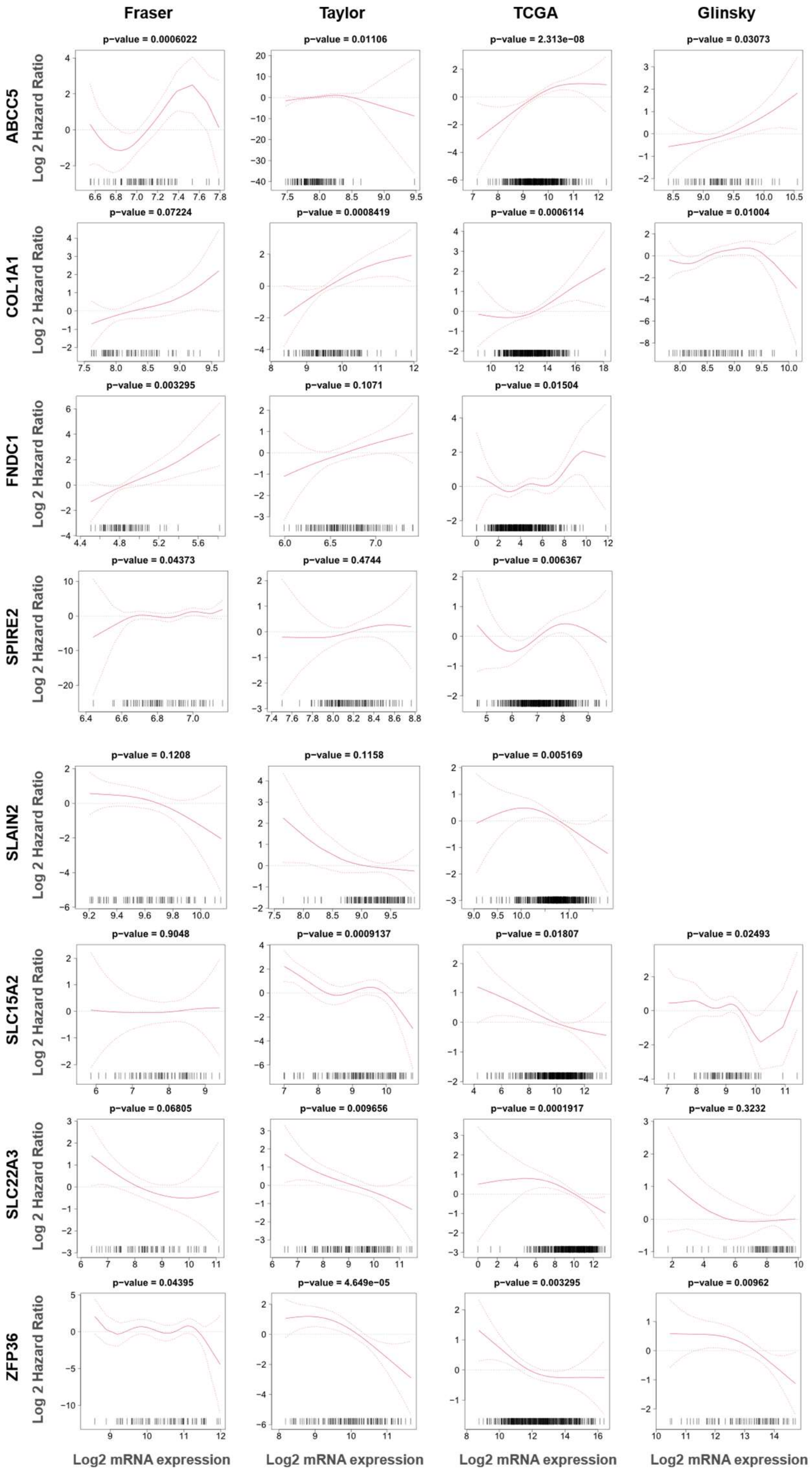


**<u>Fig S4.</u>** *Related to Fig 2*. Smooth Hazard ratio curves. x-axis represents the gene expression level and y-axis the Log Hazard ratio. The p-value indicates the significance of the association between the gene and the outcome calculated via a likelihood ratio test.

### Figure S5

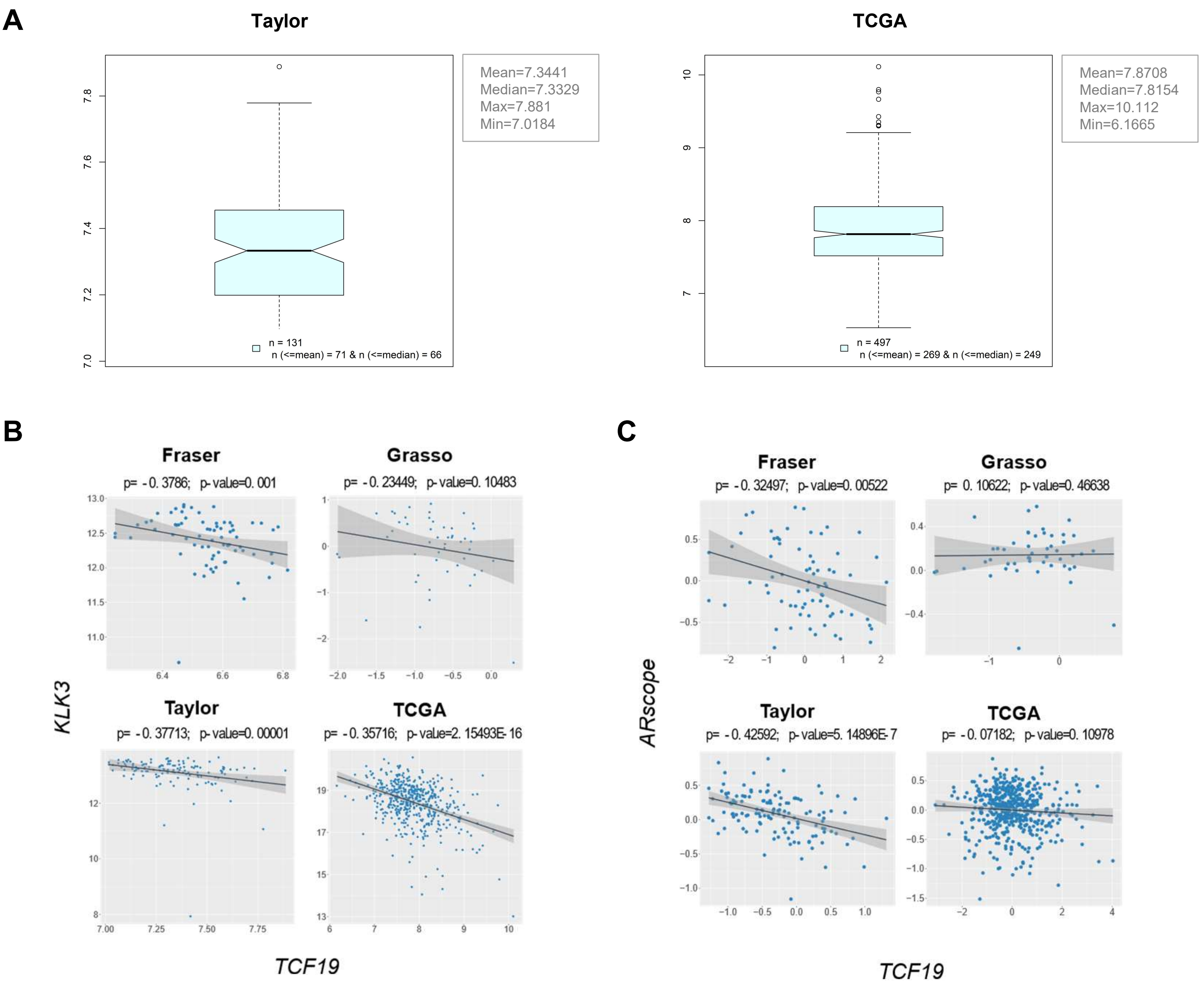


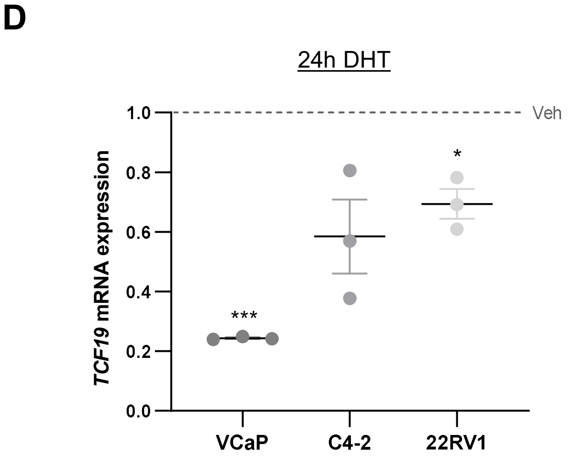


**<u>Fig S5.</u>** *Related to Fig 3.* **A)** Box plots showing the distribution of patients based on their *TCF19* expression levels. The cohort size and the mean, median, max and min values of *TCF19* expression in each indicated dataset are shown. **B and C)** Plotted values correspond to the Log$_2$-normalized gene expression values of the indicated genes or AR signature from (PMID: 17010675) (in X and Y-axis) in the primary tumor specimens from each patient in the indicated dataset. Black line represents linear regression, grey area indicates the limits of the confidence intervals and p and p-value indicate Spearman's correlation coefficient and statistical significance respectively. **D)** Analysis of *TCF19* expression by qRT-PCR in the indicated cell line treated for 24h with AR agonist (dihydrotestosterone, DHT, 10 nM). Data were normalized to *GAPDH* expression and untreated (Veh; vehicle) condition. The dotted line represents the normalized value of the vehicle data. A one sample *t*-test was performed. *n* = 3 independent experiments. *p, p-value. ns p≥0.05, * p<0.05, ** p<0.01, *** p<0.001.*

#### Figure S6

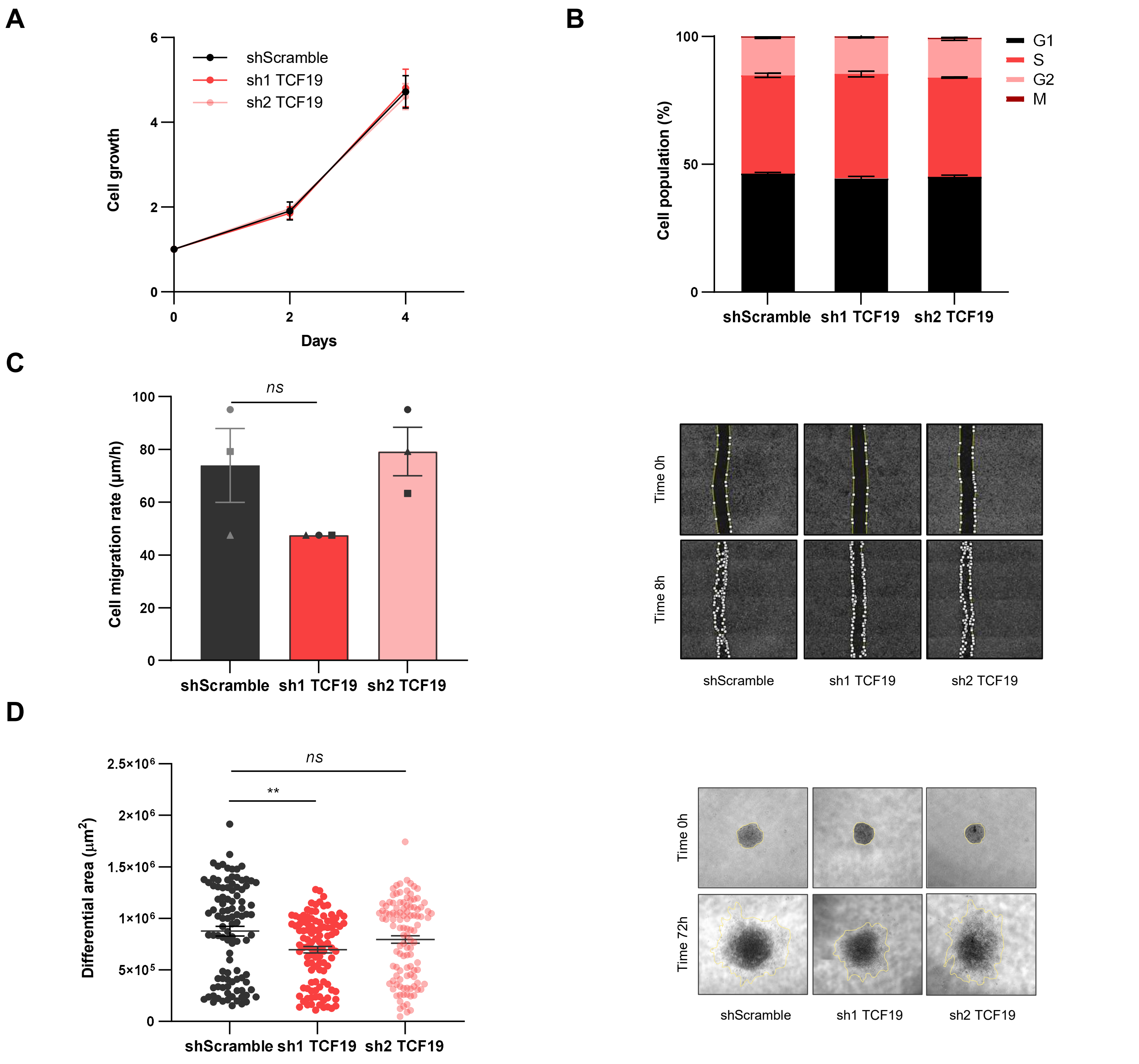


<u>Fig S6.</u> *Related to Fig 4*. **A)** Analysis of cell proliferation of PC3 cells upon *TCF19* silencing. The cell number at each time point relative to time 0 are represented. A multiple paired *t*-test was applied for statistical analysis. **B)** Cell cycle distribution of shScramble or shTCF19 transduced PC3 cells. A multiple paired *t*-test was applied for statistical analysis. **C)** Analysis of cell migration rate of PC3 cells transduced with the indicated shRNA. The different biological replicates are indicated with unique dot shapes (left panels). A two-tailed paired Student's *t*-test was applied for statistical analysis. *n*=3 independent experiments. Representative images of the scratch at initial and final timepoint are shown (right panels). **D)** Analysis of invasive growth of PC3 cells transduced with the indicated shRNA. Cell spheroids were embedded in collagen and measured at 0 and 72 hours (h). The differential area between final and initial timepoint was measured (left panel). A two-tailed unpaired Student's *t*-test was applied for statistical analysis. *n*=6, independent analysis. Representative images of the spheroids at final timepoint are shown (right panel). *p, p-value. ns* $p \geq 0.05$, * $p<0.05$, ** $p<0.01$, *** $p<0.001$.

### Figure S7

**A**

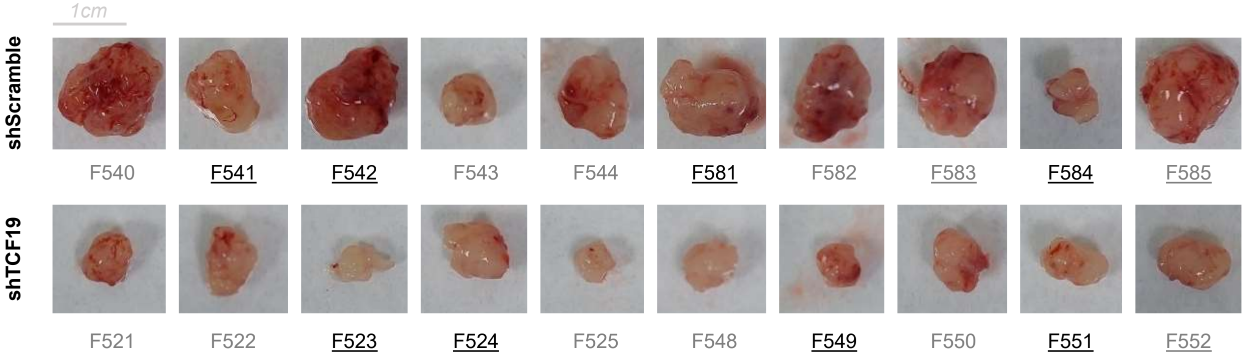


**B**

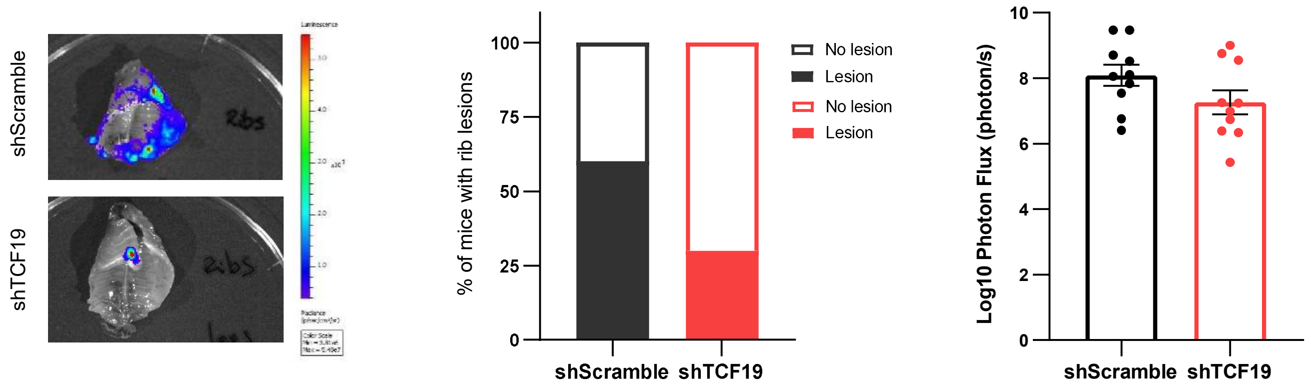


**Fig S7.** *Related to Fig 5.* **A)** Images of the tumours collected at the end of the orthotopic xenotransplant assay. The tumors characterized in Figure 6 are indicated underline (qPCR) and grey (IHC/IF). **B)** Evaluation of metastatic lesions in the ribs by orthotopic xenotransplant assay. Representative images (left panel). The *ex vivo* incidence of rib lesions (middle panel). Luciferase signal above day 0 was considered metastasis-positive. The $Log_{10}$ photon flux signal of the ribs are represented (right panel). A two-sided Fisher's exact test was performed for statistical analysis. *p, p-value. ns p≥0.05, * p<0.05, ** p<0.01, *** p<0.001.*

### Figure S8

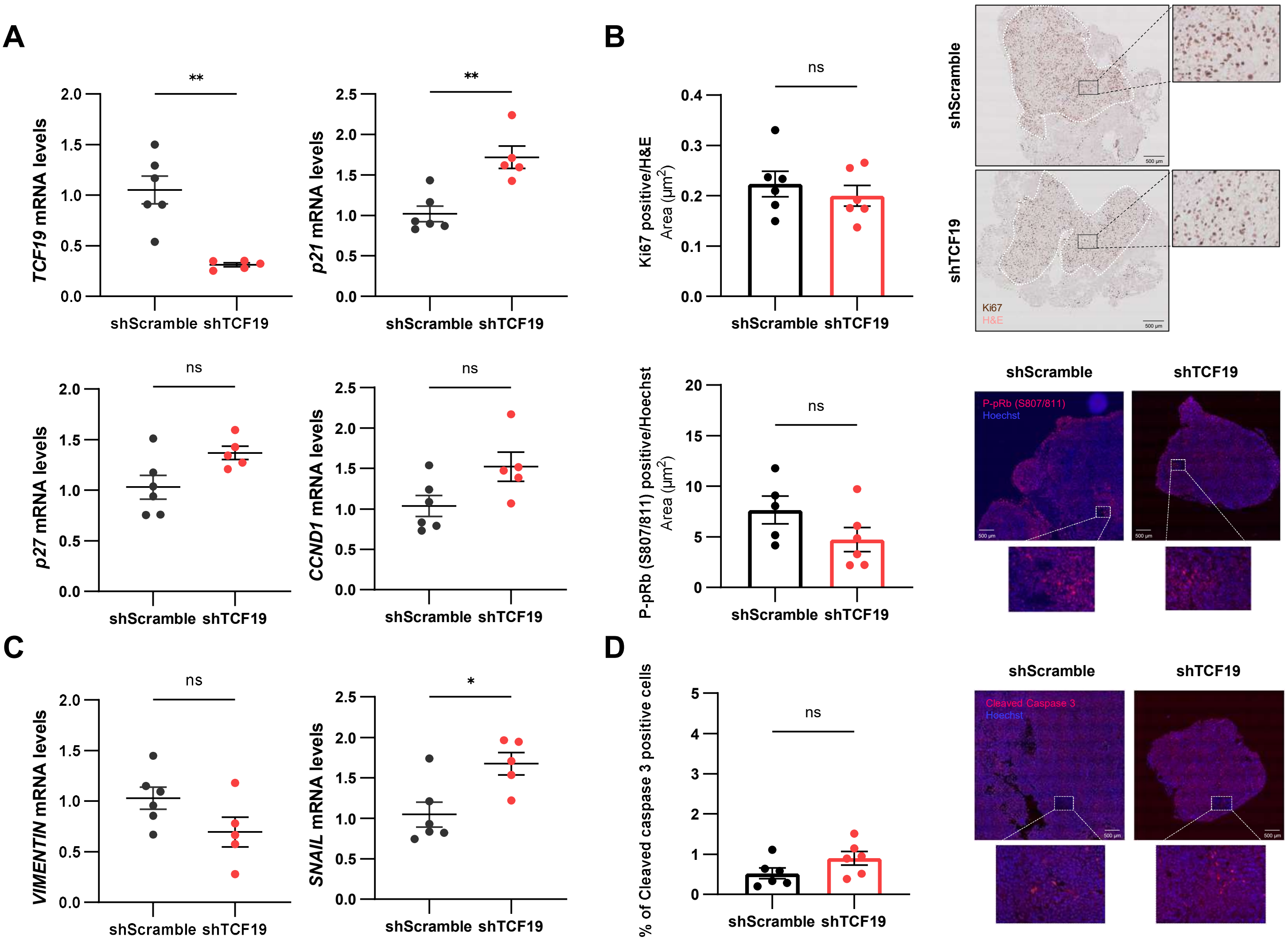


**<u>Fig S8.</u>** *Related to Fig 6*. **A)** Gene expression analysis of shScramble and shTCF19 tumors from the *in vivo* orthotopic experiment. The tumors selected for the qRT-PCR analysis are indicated in Fig S7. Data were normalized to *GAPDH* expression and shScramble condition. A Mann-Whitney *U*-test was performed for statistical analysis. **B)** Analysis of Ki67 and P-pRb (S807/811) in shScramble and shTCF19 tumors from the *in vivo* orthotopic experiment. The tumors selected for the IHC/IF analysis are indicated in Fig S7. A Mann-Whitney *U*-test was performed for statistical analysis (left panels). Representative images are shown (right panels). ***C)*** Gene expression analysis of shScramble and shTCF19 tumors from the *in vivo* orthotopic experiment. The tumors selected for the qRT-PCR analysis are indicated in Fig S7. Data were normalized to *GAPDH* expression and shScramble condition. A Mann-Whitney *U*-test was performed for statistical analysis. **D)** Analysis of Cleaved Caspase 3 in shScramble and shTCF19 tumors from the *in vivo* orthotopic experiment. The tumors selected for the IF analysis are indicated in Fig S7. A Mann-Whitney *U*-test was performed for statistical analysis (left panels). Representative images are shown (right panels). *ns p≥0.05, * p<0.05, ** p<0.01, *** p<0.001.*

#### Figure S9

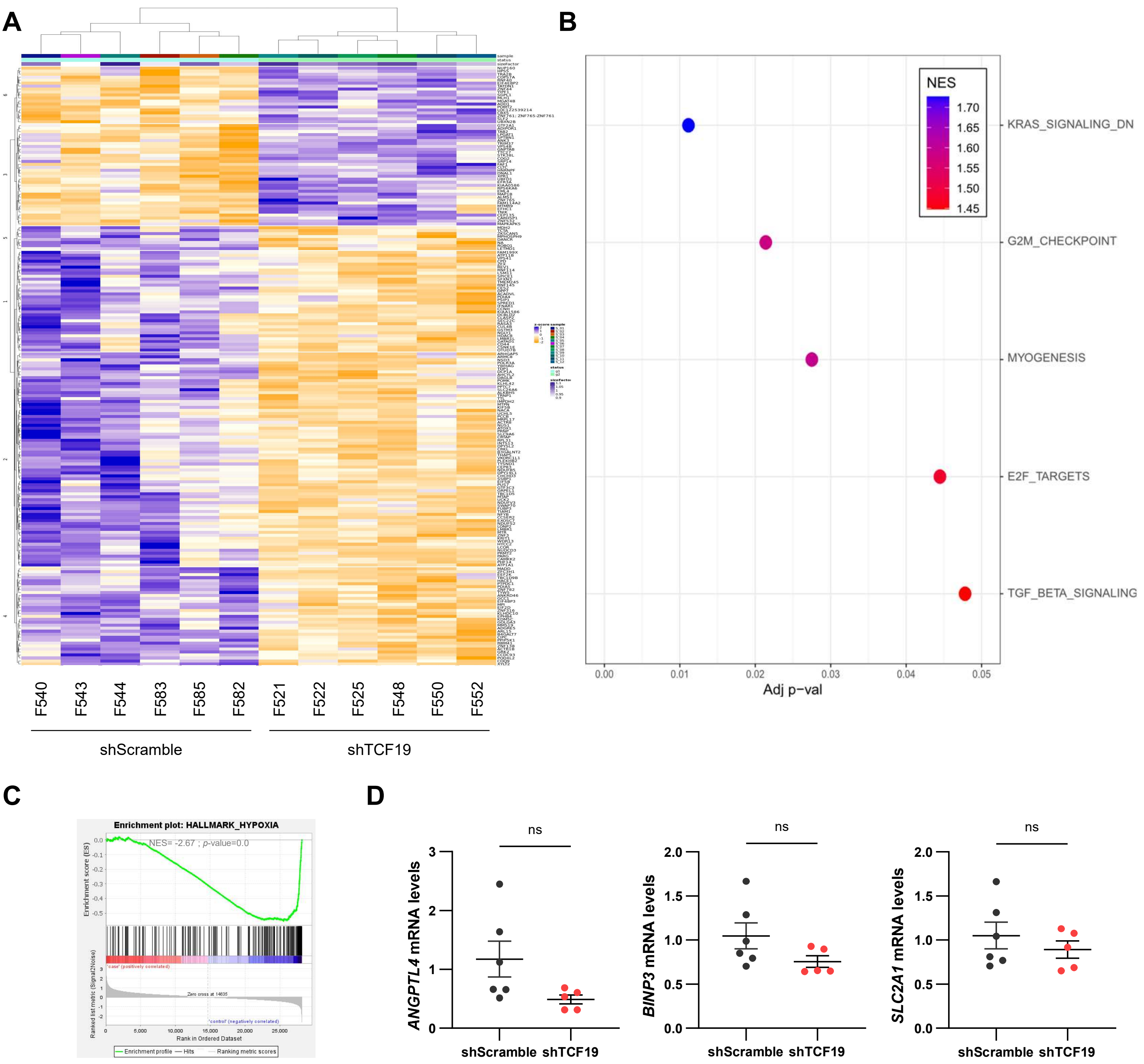


**Fig S9.** *Related to Fig 6.* **A)** The Z-scored heatmap representing the Top200 significant changes (p<0.05) upon *TCF19* depletion in the RNA-seq analysis. **B and C)** A GSEA analysis was performed to identify pathways enriched upon *TCF19* depletion. The Top 5 pathways phenotypically enriched in the shTCF19 condition at the GSEA analysis are shown (B). The GSEA profile of a selected pathway from Fig. 6B is shown (C). **D)** Gene expression analysis of shScramble and shTCF19 tumors from the *in vivo* orthotopic experiment. The tumors selected for the qRT-PCR analysis are indicated in Fig S7. Data were normalized to *RPLP0* expression and shScramble condition. A Mann-Whitney *U*-test was performed for statistical analysis. *p, p-value. ns p≥0.05, * p<0.05, ** p<0.01, *** p<0.001.*

### Figure S10

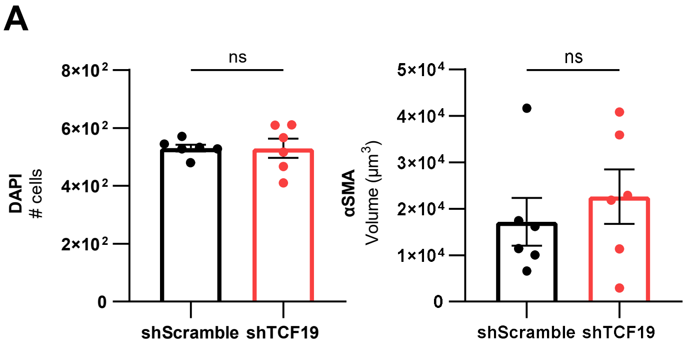


**<u>Fig S10.</u>** *Related to Fig 7*. Analysis of total αSAM and DAPI in shScramble and shTCF19 tumors from the *in vivo* orthotopic experiment. The tumors selected for the IHC/IF analysis are indicated in Fig S7. A Nested *t*-test was performed for statistical analysis. *p, p-value. ns* $p \geq 0.05$, * $p<0.05$, ** $p<0.01$, *** $p<0.001$.
